# Molecular characterization of *required for killing-2*: a gene required for spore killing by *Neurospora Sk-3*

**DOI:** 10.64898/2026.09.07.749943

**Authors:** Sabiha Sultana, Shahriar Mahmud, Parham Jazireian, Jessica M. Lohmar, Daren Brown, Thomas M. Hammond, Nicholas A. Rhoades

## Abstract

The *Sk-3* selfish genetic element in *Neurospora* fungi is transmitted to offspring in a biased manner through spore killing. In crosses between an *Sk-3* strain and a *Spore killing-sensitive* (*Sk-S*) strain, nearly all offspring that fail to inherit *Sk-3* are killed. Although a mutation called *rfk-2*^UV^, which disrupts spore killing by *Sk-3*, was isolated in an earlier work, its precise location within the *Sk-3* element was not identified. Here, we demonstrate that *rfk-2*^UV^ is a T>G transversion mutation within a protein coding gene called *rfk-2*. The *rfk-2* gene is located on Chromosome III, and it encodes a transcript that undergoes RNA splicing and at least two RNA editing events. One of these editing events changes a UAG stop codon to a UGG tryptophan codon and allows for production of a 102 amino acid protein. Additionally, while our results demonstrate that *rfk-2* is required for spore killing, they also show that it is insufficient for the process. *Sk-3* is thus more complex than previously anticipated. Rather than using a single gene to produce its killer, *Sk-3* requires multiple genes to kill spores and break the rules of Mendelian Inheritance.

## INTRODUCTION

In sexually reproducing organisms, most genetic elements are fairly transmitted from parents to offspring according to rules discovered by Gregor Mendel in the 19^th^ century. Some genetic elements, however, are transmitted from parents to offspring in a biased manner. Well known examples are meiotic drive elements and gametic drive elements. Meiotic drive elements gain transmission advantage during meiosis. For example. the *Ab10* element in maize achieves drive by manipulating chromosome segregation so that it has a better than even chance of being placed in an egg cell over a polar body (Dawe 2022). Gametic drive elements work after meiosis, and they typically achieve drive by pitting gametes against one another. *Segregation Distorter* in *D. melanogaster* is an example of a gametic drive element that directs *Sd* carrying-sperm to target those lacking *Sd* for destruction (Larracuente and Presgraves 2012).

Recent work has revealed that gametic drive elements are widespread in fungi, where they are commonly referred to as killer meiotic drivers (KMDs; Zanders and Johannesson 2021). Three of the first examples of KMDs were discovered by Turner and colleagues in Neurospora fungi. These KMDs were named *Spore killer-1* (*Sk-1*), *Spore killer-2* (*Sk-2*), and *Spore killer-3* (*Sk-3*), and as their names imply, they gain transmission advantage through spore killing (Raju 1979; Turner and Perkins 1979). *Sk-1* was found in *N. sitophila* and it appears to encode all of the information it needs for its biased transmission mechanism within a single gene (Svedberg et al. 2021). *Sk-2* and *Sk-3*, which were identified in *N. intermedia* and introgressed into *N. crassa* for genetic analysis, are complex KMDs that span millions of base pairs and contain hundreds of genes (Campbell and Turner 1987; Svedberg et al. 2018).

Spore killing is relatively easy to detect within asci (spore sacs) of the model Neurospora species *N. crassa.* In this organism, ascus development begins with the fusion of two haploid nuclei, one from each parent of a cross (Raju 1980). The diploid nucleus then proceeds through meiosis to produce four nuclei, each of which undergoes a single round of mitosis. The resulting eight nuclei are then incorporated into developing ascospores, one nucleus per ascospore. Assuming each ascospore completes development without complication, the ascus should contain eight black (melanized) ascospores at maturity. Indeed, this is the expected phenotype of asci from a cross between two non-spore killer strains. In contrast, when a cross is performed between a Spore killer strain such as *Sk-2* or *Sk-3* and a *Spore killing sensitive* strain (*Sk-S*), mature asci contain four black ascospores and four shriveled, white (hyaline) ascospores. Only the black ascospores are viable, and they almost always have the spore killer genotype (>99%, Turner and Perkins 1979).

*Sk-2* and *Sk-3* are located within a similar centromere spanning interval of Chromosome III, and they are both transmitted to offspring as single haplotypes due to recombination suppression (Campbell and Turner 1987). Although the mechanism of recombination suppression has not been tested directly, it is thought to result from a series of unique inversions that exist within each element relative to each other and the corresponding Chromosome III interval in *Sk-S* strains (Svedberg et al. 2018). Recombination suppression likely serves to keep the genes controlling *Sk-2* and *Sk-3*’s biased transmission from separating during meiosis, and a "killer neutralization" model has been proposed to explain how this biased transmission is achieved (Hammond et al. 2012). This model holds that both Spore killers encode a poison (a killer protein) and an antidote (a resistance protein), and while both the poison and antidote are present throughout the ascus during early stages of development, the antidote becomes restricted to those ascospores that encode it during late stages of development, and without access to the antidote, *Sk-S* ascospores succumb to the poison. The killer neutralization model is supported by the discovery of an antidote that provides resistance to *Sk-2* and *Sk-3* spore killing. The antidote is encoded by the *rsk* gene, and the type of resistance provided by the gene depends on its allele (Hammond et al. 2012). For example, the *rsk* allele in *Sk-2* provides resistance to the *Sk-2* killer and the *rsk* allele in *Sk-3* provides resistance to the *Sk-3* killer. The killer neutralization model is also supported by the discovery of a poison protein called RFK-1 (Harvey et al. 2014; Rhoades et al. 2019). This protein, encoded by the *rfk-1* gene, is required for spore killing by *Sk-2*. Interestingly, sequence analysis of the *Sk-3* genome has failed to identify an obvious RFK-1 homolog (Svedberg et al. 2018), suggesting that *Sk-2* and *Sk-3* may use unrelated proteins as their poisons.

In a previous study, we identified a mutation that disrupts spore killing by *Sk-3* (Velazquez et al. 2022). We named this mutation *rfk-2*^UV^ and mapped it to the left arm of Chromosome III. Here, we identify and molecularly characterize the *rfk-2* gene. While *rfk-2* is indeed required for spore killing by *Sk-3*, our results demonstrate it to be insufficient for the process. The implications of our findings with respect to the evolutionary origins and spore killing mechanisms of *Sk-2* and *Sk-3* are discussed.

## MATERIALS AND METHODS

### Neurospora strains, media, crosses, and culture conditions

Strain genotypes are provided in Table 1 and strain derivation information is provided in Table S1. Vogel’s minimal medium (VMM) or Vogel’s minimal agar (VMA) containing 2% agar was used for vegetative propagation of all strains (Vogel 1956). Hygromycin B (GoldBio, H-270) and G418 sulfate (GoldBio, G-418) were used at 200 μg per ml and 900 μg per ml, respectively, to select for resistance to each antibiotic. Cultures were incubated at 32 °C in an incubator or at room temperature on laboratory shelves. Synthetic Crossing Agar (SCA; 1.5% sucrose, 6.5 pH, 2% Agar) was used for crosses (Westergaard and Mitchell 1947). Crosses were performed at room temperature. All strains used in this study carried a mutant *rid* allele to suppress the genome defense process of Repeat-Induced Point Mutation (Freitag et al. 2002). The *fl* mutation was used to prevent macroconidiation by strains used as designated females in unidirectional crosses (Perkins et al. 2000). Mutant alleles of *mus-51* were used to improve transformation efficiency (Ninomiya et al. 2004). Unless otherwise indicated, at least one parent of each cross used in a spore killing assay or a gene drive assay carried a *sad-2*^Δ^ allele, which suppresses Meiotic Silencing by Unpaired DNA and allows for the expression of transgenes during meiosis (Shiu et al. 2006).

**Table 1.** Strains used in this study.

| Strain name | Genotype |
| --- | --- |
| F2-23 | *rid; fl; Sk-S A<sup>+</sup>* |
| F2-26 | *rid; fl; Sk-S a<sup>+</sup>* |
| FGSC-10340 | *rid; Sk-S; mus-51<sup>RIP70</sup> a<sup>+</sup>* |
| ISU-3036 | *rid; fl; Sk-S; sad-2<sup>Δ</sup>::hph<sup>+</sup> A<sup>+</sup>* |
| ISU-3037 | *rid; fl; Sk-S; sad-2<sup>Δ</sup>::hph<sup>+</sup> a<sup>+</sup>* |
| ISU-3291 | *rid; Sk-3; mus-51<sup>Δ</sup>::bar<sup>+</sup> A<sup>+</sup>* |
| ISU-4677 | *rid; Sk-3 rsk<sup>Δ</sup>::hph<sup>+</sup>; mus-51<sup>Δ</sup>::bar<sup>+</sup> A<sup>+</sup>* |
| ISU-4683 | *rid; Sk-3 rfk-2<sup>UV</sup> a<sup>+</sup>* |
| ISU-4684 | *rid; Sk-3 rfk-2<sup>UV</sup>; mus-51<sup>Δ</sup>::bar<sup>+</sup> a<sup>+</sup>* |
| ISU-4773 | *rid; Sk-3 rsk<sup>Δ</sup>::hph<sup>+</sup> rfk-2<sup>UV</sup>; mus-51<sup>Δ</sup>::bar a<sup>+</sup>* |
| ISU-5211 | *rid; Sk-3 i383<sup>Δ</sup>::hph<sup>+</sup> A<sup>+</sup>* |
| ISU-5295 | *rid; Sk-3 i388<sup>Δ</sup>::hph<sup>+</sup> a<sup>+</sup>* |
| ISU-5331 | *rid; Sk-3 rfk-2<sup>UV</sup>; mus-51<sup>Δ</sup>::bar a<sup>+</sup>* |
| ISU-5332 | *rid; Sk-3 rfk-2<sup>UV</sup>; mus-51<sup>Δ</sup>::bar a<sup>+</sup>* |
| ISU-5333 | *rid; Sk-3 rfk-2<sup>UV</sup>; mus-51<sup>Δ</sup>::bar a<sup>+</sup>* |
| ISU-5334 | *rid; Sk-3 rfk-2<sup>UV</sup>; mus-51<sup>Δ</sup>::bar a<sup>+</sup>* |
| ISU-5335 | *rid; Sk-3 rfk-2<sup>UV</sup>; mus-51<sup>Δ</sup>::bar a<sup>+</sup>* |
| ISU-5337 | *rid; Sk-3 i350<sup>Δ</sup>::hph<sup>+</sup>; mus-51<sup>Δ</sup>::bar<sup>+</sup> A<sup>+</sup>* |
| ISU-5338 | *rid his-3<sup>+</sup>::iSPS7(rfk-2<sup>+</sup>)-hph<sup>+</sup>; Sk-S; mus-51<sup>RIP70</sup> a<sup>+</sup>* |
| ISU-5434 | *rid; Sk-3 i358<sup>Δ</sup>::hph<sup>+</sup>; mus-51<sup>Δ</sup>::bar<sup>+</sup> A<sup>+</sup>* |
| ISU-5462 | *rid; Sk-3 i397<sup>Δ</sup>::hph<sup>+</sup> a<sup>+</sup>* |
| ISU-5563 | *rid; fl; sad-2<sup>Δ</sup>::genR<sup>+</sup> A<sup>+</sup>* |
| ISU-5564 | *rid; fl; sad-2<sup>Δ</sup>::genR<sup>+</sup> a<sup>+</sup>* |
| ISU-5567 | *rid; Sk-3 i397<sup>+</sup>/i397<sup>Δ</sup>::hph<sup>+</sup> rfk-2<sup>UV</sup>/?; mus-51<sup>Δ</sup>::bar<sup>+</sup> a<sup>+</sup>* |
| ISU-5568 | *rid; Sk-3 i397<sup>+</sup>/i397<sup>Δ</sup>::hph<sup>+</sup> rfk-2<sup>UV</sup>; mus-51<sup>Δ</sup>::bar<sup>+</sup> a<sup>+</sup>* |
| ISU-5569 | *rid; Sk-3 i397<sup>+</sup>/i397<sup>Δ</sup>::hph<sup>+</sup> rfk-2<sup>UV</sup>/rfk-2<sup>+</sup>; mus-51<sup>Δ</sup>::bar<sup>+</sup> a<sup>+</sup>* |
| ISU-5571 | *rid; Sk-3 i397<sup>+</sup>/i397<sup>Δ</sup>::hph<sup>+</sup> rfk-2<sup>UV</sup>; mus-51<sup>Δ</sup>::bar<sup>+</sup> a<sup>+</sup>* |
| ISU-5846 | *rid his-3<sup>+</sup>::iSPS7(rfk-2<sup>+</sup>)-hph<sup>+</sup>; Sk-S; mus-51<sup>RIP70</sup> a<sup>+</sup>* |
| ISU-5848 | *rid; Sk-3 i383<sup>Δ</sup>::hph<sup>+</sup> mus-51<sup>RIP70</sup> A<sup>+</sup>* |
| ISU-5852 | *rid his-3<sup>+</sup>::iSPS7(rfk-2<sup>+</sup>)-hph<sup>+</sup>; Sk-3 i383<sup>Δ</sup>::hph<sup>+</sup>; a<sup>+</sup>* |
| ISU-5856 | *rid; Sk-3 i397<sup>Δ</sup>::hph<sup>+</sup> rfk-2<sup>UV</sup>; mus-51<sup>Δ</sup>::bar<sup>+</sup> a<sup>+</sup>* |
| ISU-5859 | *rid; Sk-3 i397<sup>Δ</sup>::hph<sup>+</sup>; mus-51<sup>Δ</sup>::bar<sup>+</sup> a<sup>+</sup>* |
| P6-07 | *rid A<sup>+</sup>* |

### *Sk-3* reference sequence and naming conventions

All fungal strains used in this study are *N. crassa* isolates. The *Sk-3* reference sequence, however, was obtained from the genome of *N. intermedia Sk-3* strain FGSC 3194 (Svedberg et al. 2018). The *rfk-2*^UV^ mutation was isolated in an *N. crassa Sk-3* strain and it was subsequently mapped centromere-proximal of a short DNA interval called *i322*. This short (2 bp) interval corresponds to Chromosome III, Positions 1,017,772–1,017,773, of the *Sk-3* reference sequence (Velazquez et al. 2022). Typically, the DNA intervals mentioned in this study are named after the transformation vector used to delete or modify each interval. For example, Transformation Vector v350 deletes DNA interval *i350* and replaces it with a hygromycin resistance cassette (*hph*).

### SNV identification

Genomic DNA isolations were performed with the IBI Scientific Mini Genomic DNA Isolation Kit for Plants and Fungi (IB47231) or the Zymo Research Quick-DNA Fungal/Bacterial Miniprep Kit. DNA from six *Sk-3 rfk-2*^UV^ strains was combined in equal ratios for whole genome sequencing as follows: DNA from ISU-5331 and ISU-5332 was mixed for Dataset A (NCBI SRA: SRX34189115); DNA from ISU-5333 and ISU-5334 was mixed for Dataset B (NCBI SRA: SRX34189116); and DNA from ISU-5335 and ISU-4684 was mixed for Dataset C (NCBI: SRA SRX34189117). Datasets A–C were combined to produce the *rfk-2*^UV^ sequencing read dataset. DNA from one *Sk-3 rfk-2^+^* strain (ISU-3291) was subjected to whole genome sequencing twice to produce Dataset D (NCBI SRA: SRX34189118) and Dataset E (NCBI SRA: SRX34189119). Datasets D and E were combined to produce the *rfk-2^+^* sequencing read dataset. DNA libraries were prepared and sequenced, and raw sequencing reads were processed, as previously described (Lohmar et al. 2022). The *rfk-2*^UV^ and *rfk-2^+^* datasets were then mapped to Chromosome III, Positions 1 to 1,017,702, of the *Sk-3* reference sequence with Bowtie2 (Langmead and Salzberg 2012). Samtools mpileup was used to produce *rfk-2*^UV^ and *rfk-2*^+^ pileup text files (Li et al. 2009). Samtools mpileup settings were as follows: the minimum base quality score was set to zero and the read-pair overlap detection process, the anomalous read pair discarding mechanism, as well as the per-base alignment quality function were disabled. Variant positions in *rfk-2*^UV^ were called from the *rfk-2*^UV^ pileup document with a custom Python script. Positions that were covered by more than 10 reads with more than 90% of the mapped bases different from the reference sequence were defined as variants. The *rfk-2*^UV^ variant positions were then examined in the *rfk-2^+^* pileup document to determine if the variants were unique to the *rfk-2*^UV^ strains or if they were common to both the *rfk-2^+^* and *rfk-2*^UV^ strains. Galaxy (Version 26.1) was used to run Bowtie2 and Samtools (The Galaxy Community 2024).

### *N. crassa* transformation and vector construction

Transformations were performed by electroporation of macroconidia as previously described (Margolin et al. 1997; Rhoades et al. 2020). Double-Joint PCR (Yu et al. 2004) was used to construct Transformation Vectors v350, v358, v383, v388, v397, and v417. Primers are described in Table S2. Genomic DNA from ISU-3291 was used as template for the left flank and the right flank of all vectors. Plasmid pTH1119.2 (GenBank: PX241540.1) or pTH1256.1 (GenBank: MH550659.1) was used as the template for the center fragment for all vectors except v417, which used Plasmid pGENNotI (Desjardins et al. 2004; Lohmar et al. 2022). ISU-3291 and FGSC-10340 were used as the *Sk-3* and *Sk-S* transformation hosts, respectively. Plasmid pSPS5.4 was constructed by cloning a 2.4 kb PCR product obtained from genomic DNA of ISU-3291 with Primer Set 2507/2508 into pJET1.2 with the CloneJET PCR Cloning Kit (Thermo Scientific). Plasmid pSPS7.2 was constructed by transferring the 2.4 kb *Not*I fragment from pSPS5.4 to the *Not*I site of pTH1256.1. Plasmid pSPS7.2 was linearized with *Ssp*I before transformation of FGSC-10340.

### Qualitative spore killing assays and quantitative gene drive assays

Spore killing assays were performed with unidirectional crosses. The spore killing assays are qualitative because they involve imaging asci and visually assessing the presence or absence of spore killing. Each designated female strain carried the *fl* mutant allele. Designated female strains were cultured on SCA in 60 mm culture dishes for 6–10 days to allow protoperithecia to develop. Tissue suspensions (conidia and/or mycelia in water) were made for each male strain, and approximately 200 μl of suspension were placed over the surface of the female cultures in 5– 10 μl drops. Cultures were then incubated on culture shelves at room temperature with ambient lighting. To visually score spore killing, perithecia were harvested approximately 12 days post fertilization and asci were dissected from perithecia into 25% glycerol. The asci were then imaged under magnification. Quantitative gene drive assays were performed to measure the transmission rates of specific genetic elements (e.g., *Sk-3*, *rfk-2^+^*). Gene drive assays were performed similarly to the spore killing assays, except instead of dissecting perithecia to visualize asci, ascospores were collected from the undersides of the crossing lids 24 days or later after fertilization. Ascospore suspensions were then stored in the dark at 4 °C for at least 18 hours before heat shock at 60 °C for 30 minutes. Ascospores were then spread across VMA, and individual ascospores were transferred to 16 × 125 mm glass culture tubes containing VMA. The resulting cultures were then genotyped for the presence of *hph* with media-based hygromycin resistance assays. When more than one genotype could confer resistance to hygromycin, genotypes were determined by genomic DNA isolation and PCR.

### *rfk-2* transcript prediction

RNA sequencing datasets were downloaded for *Sk-S* × *Sk-3* crosses (NCBI SRA: SRR7700945, SRR7700971, and SRR7700972) and *Sk-S* × *Sk-S* crosses (NCBI SRA: SRR7700966, SRR7700967, and SRR7700968). The six datasets were combined into two datasets by cross type. HiSat2 (Kim et al. 2019) was then used with default settings to map all reads from each dataset to an *N. intermedia Sk-S* genome (FGSC 8716; Svedberg et al. 2018). Samtools view was used to export reads that mapped concordantly to the *spo-11* locus (Chromosome V: 3,216,276*–* 3,223,712) in SAM format (primary alignments only). Reads that did not align to the *N. intermedia Sk-S* genome were collected into "unmapped read" files. HiSat2 was then used to map the "unmapped reads" to the *rfk-2* locus (*Sk-3* reference, Chromosome III: Positions 332,122*–* 333,160). Samtools view was used to export all reads that mapped concordantly to the *rfk-2* locus in SAM format (primary alignments only). A custom Python script was used to determine read counts per position. Galaxy (Version 26.1) was used to run HiSat2 and Samtools (The Galaxy Community 2024).

### RNA isolation and cDNA cloning

Crosses were performed on 18.0 ml SCA in 60 mm culture dishes at room temperature. Designated female strains were cultured for six days, after which they were fertilized at three locations with three 33 μl aliquots of conidia in water (1000 conidia per μl) from a designated male strain. Perithecia were harvested from crossing plates eight days after fertilization and ground in liquid nitrogen with a mortar and pestle. A liquid nitrogen chilled microspatula was then used to transfer approximately 100 mg of frozen ground tissue into 1.2 ml of TRIzol (Thermo Scientific) before extracting total RNA according to the manufacturer’s recommended protocol. A DNAse I treatment step was performed for some samples (specified below), followed by purification with the Monarch Spin RNA Cleanup Kit (New England BioLabs). The ProtoScript II First Strand cDNA Synthesis Kit (New England BioLabs) with the provided "d(T)23 VN" primer was used to synthesize cDNA according to the manufacturer’s recommended protocol.

## RESULTS

### DNA interval *i350* is required for spore killing

To refine the location of *rfk-2*^UV^, we sequenced the genomes of an *rfk-2^+^* strain and six descendants of the original *rfk-2*^UV^ strain (Table 1). We then mapped the resulting reads to an *Sk-3* reference sequence and identified 41 variant positions (SNVs) that were unique to *rfk-2*^UV^ genotypes (Figure 1A, Table S3). To determine which, if any, of these *rfk-2*^UV^-specific SNVs disrupted spore killing, we deleted a single DNA interval called *i350* from an *rfk-2^+^* strain with the intention of using the resulting *i350*^Δ^ allele and the *rfk-2*^UV^ SNVs to refine the position of *rfk-2* through recombination mapping. However, we found that deleting *i350* from an *rfk-2*^+^ strain eliminated the strain’s ability to kill spores (Figure 1C). This somewhat fortuitous discovery suggested that *rfk-2* may be located within associated with *i350*.

**Figure 1.**
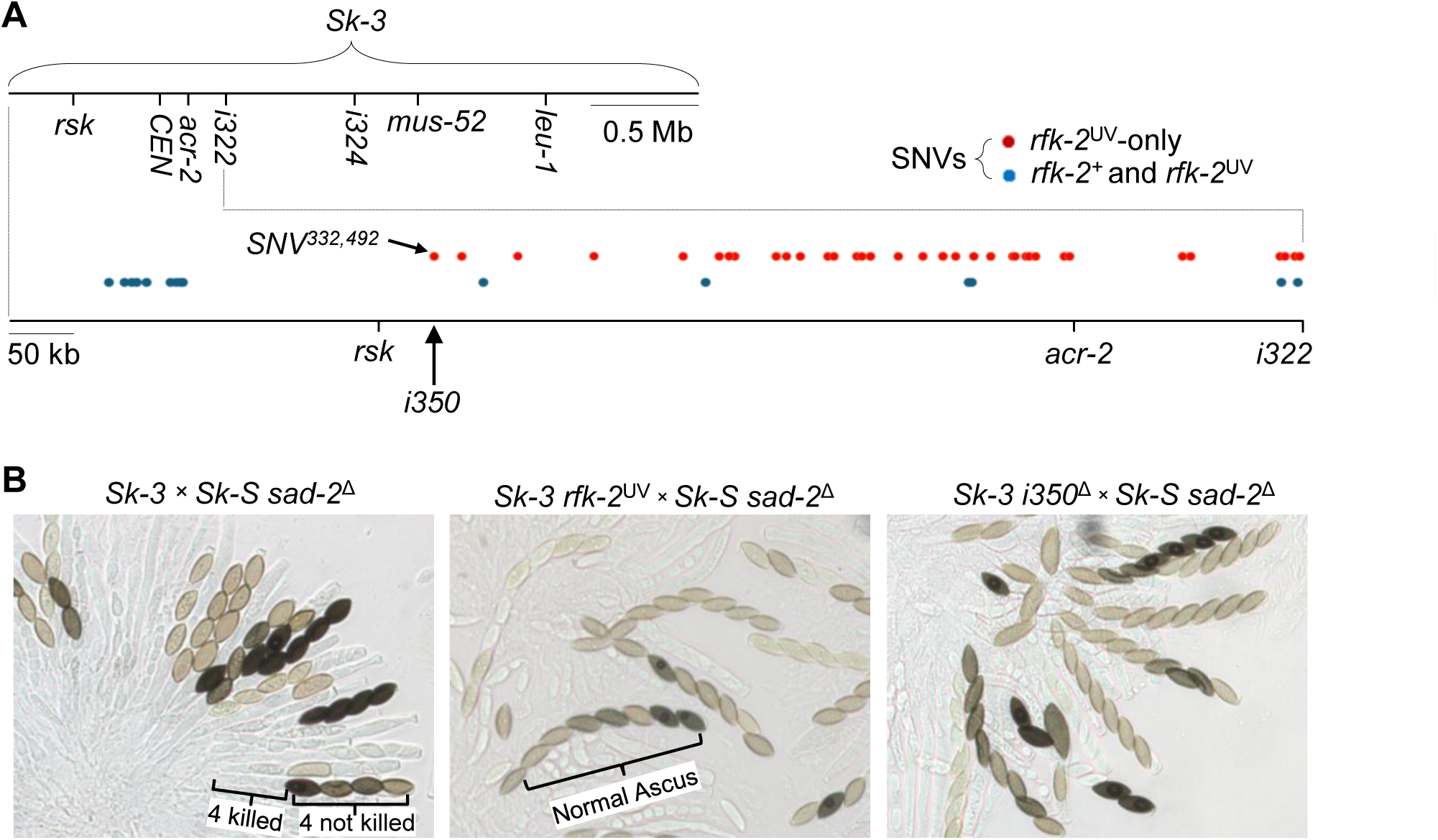
Identification of *i350*, a DNA interval required for spore killing. (A) Top of panel: A diagram of the >3 Mb *Sk-3* reference sequence is shown along with the locations of *rsk, CenIII, acr-2, i322, i324, mus-52,* and *leu-1.* Previous work mapped *rfk-2*^UV^ to the left of *i322.* Bottom of panel: The locations of SNVs between the left end of Chromosome III and *i322* are shown. SNVs with blue dots are common to the sequenced *rfk-2^+^* (ISU-3291) and *rfk-2*^UV^ strains (ISU-5531 through ISU-5535, ISU-4684). SNVs that are unique to the sequenced *rfk-2^UV^* strains are shown in red. These are assumed to be mutations that were induced during the mutagenesis screen that created *rfk-2*^UV^. (B) Deletion of *i350* disrupts spore killing. Asci from the following crosses are shown: *Sk-3* (ISU-3291) × *Sk-S sad-2*^Δ^ (ISU-3037), *Sk-3 rfk-2*^UV^ (ISU-4684) × *Sk-S sad-2*^Δ^ (ISU-3036), and *Sk-3 i350*^Δ^ (ISU-5337) × *Sk-S sad-2*^Δ^ (ISU-3037). The *Sk-3* × *Sk-S sad-2*^Δ^ cross produces asci with four black ascospores and four shriveled/hyaline ascospores. The other two crosses produce asci with eight black ascospores. Asci develop asynchronously in *N. crassa* and pigment level increases as asci develop. Ascospores with any pigment (light tan to dark black) have escaped spore killing.

### *rfk-2*^UV^ is a T>G point mutation

The *i350* interval spans a single *rfk-2*^UV^ SNV (Figure 1A; Position 332,492) and our sequencing data indicated that the position should encode a "T" in r*fk-2*^+^ strains and a "G" in *rfk-2*^UV^ strains (Table 3). To confirm this hypothesis, we isolated genomic DNA from the *rfk-2^+^*strain that was mutated to produce *rfk-2*^UV^ and the original *rfk-2*^UV^ isolate. We then used PCR to obtain amplicons from each genomic DNA sample (Figure 2A). Subsequent cloning and sequencing of the amplicons confirmed that the *rfk-2*^+^ and *rfk-2*^UV^ strains contain thymidine and guanosine nucleotides at Position 332,492, respectively (Figure 2, B and C). Next, to test if the T>G mutation is the reason why *rfk-2*^UV^ disrupts spore killing, we attempted to repair the mutation in an *rfk-2*^UV^ strain. The repair strategy involved inserting a hygromycin resistance cassette (*hph*) approximately 370 bp away (centromere-distal) from the mutation site (Figure 3A). The centromere-proximal recombination flank of the transformation vector overlaps the mutation site. Repair of the mutation results when this recombination flank exchanges the mutated nucleotide at the target site for the correct nucleotide in the recombination flank (Figure 3A, a crossover in Interval 2 leads to repair, a crossover in Interval 1 does not). We recovered four transformants and detected successful repair in one of them. We then examined the ability of one unrepaired transformant and the repaired transformant to kill spores. As expected, the unrepaired transformant produced normal asci without evidence of spore killing. In contrast, the repaired transformant produced asci with a spore killing phenotype and *Sk-3* was transmitted to 50 out of 50 offspring (Figure 3D; Chi square, p < 0.0001). These results demonstrate that repairing the T>G mutation at Position 332,492 in an *rfk-2*^UV^ strain restores spore killing.

**Figure 2.**
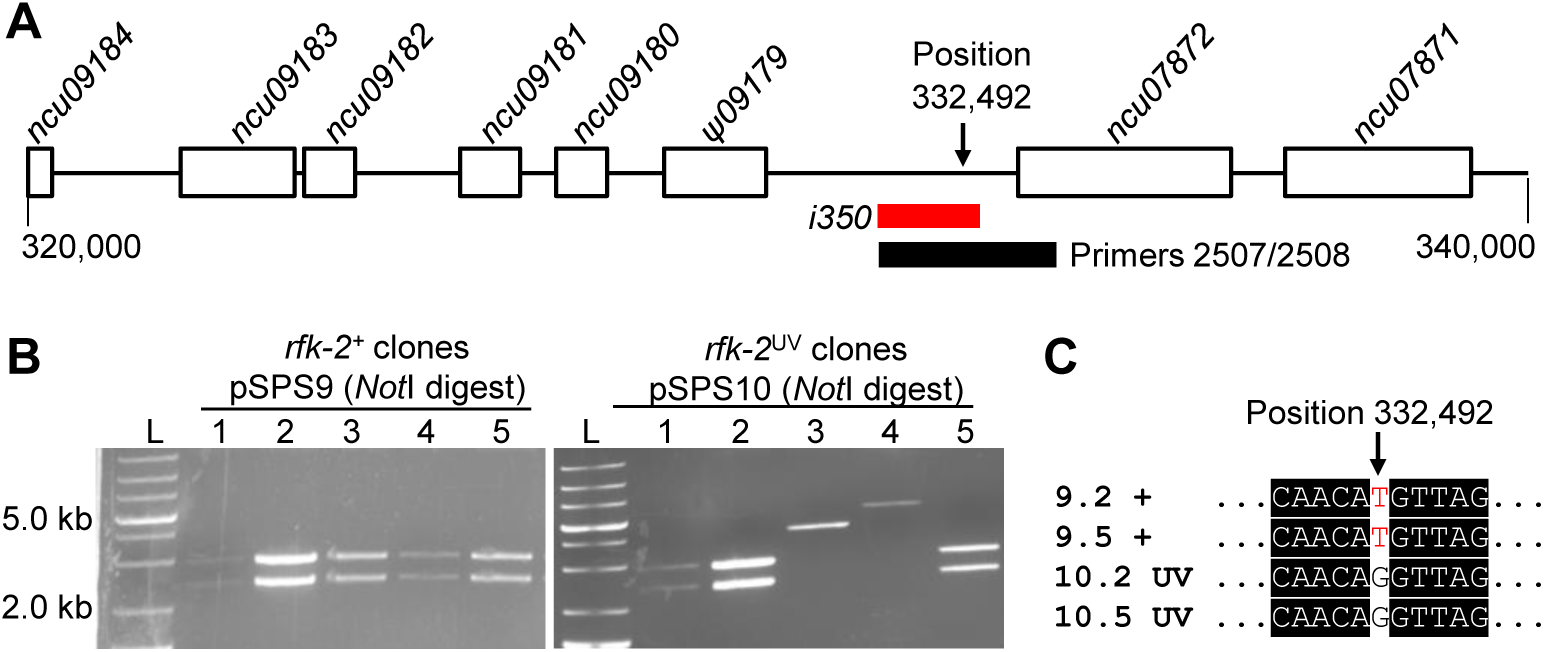
DNA sequencing of Position 332,492 in *rfk-2^+^* and *rfk-2*^UV^ strains. (A) A map of Positions 320,000 through 340,000 of Chromosome III in the *Sk-3* reference sequence is shown. Protein-coding genes were identified by matching sequences to a database of predicted *N. crassa* proteins (Galagan et al. 2003; Basenko et al. 2018). Predicted coding regions for each gene are shown as open rectangles. Genes are named according to their closest match in the *N. crassa* database. One gene appears to have undergone pseudogenization (ψ*09179).* The location of *i350* is denoted with a red bar. Primer Set 2507/2508 amplifies the region marked with a black bar. The diagram was drawn to scale with the red bar equaling 1,320 bp. (B) PCR was used to generate and clone a 2507/2508 amplicon from the original *rfk-2*^UV^ isolate (ISU-4773) and its *rfk-2^+^* progenitor (ISU-4677). (B) The amplicons were cloned into pJET1.2 with the CloneJET PCR Cloning Kit. Cloned amplicons were screened by restriction digestion and gel electrophoresis with ethidium bromide staining. Clones pSPS9.2 and pSPS9.5, with the *rfk-2^+^* PCR amplicon, and pSPS10.2 and pSPS10.5, with *rfk-2*^UV^ amplicon, were selected for whole plasmid sequencing (Plasmidsaurus Inc.). (C) Sequences were aligned with ClustalW in BioEdit (Hall 1999). An 11 bp interval of the sequence alignment is shown. The four sequences were identical except for a thymidine at Position 332,492 in the *rfk-2^+^*-clones and a guanosine at the same position in the *rfk-2*^UV^-clones.

**Figure 3.**
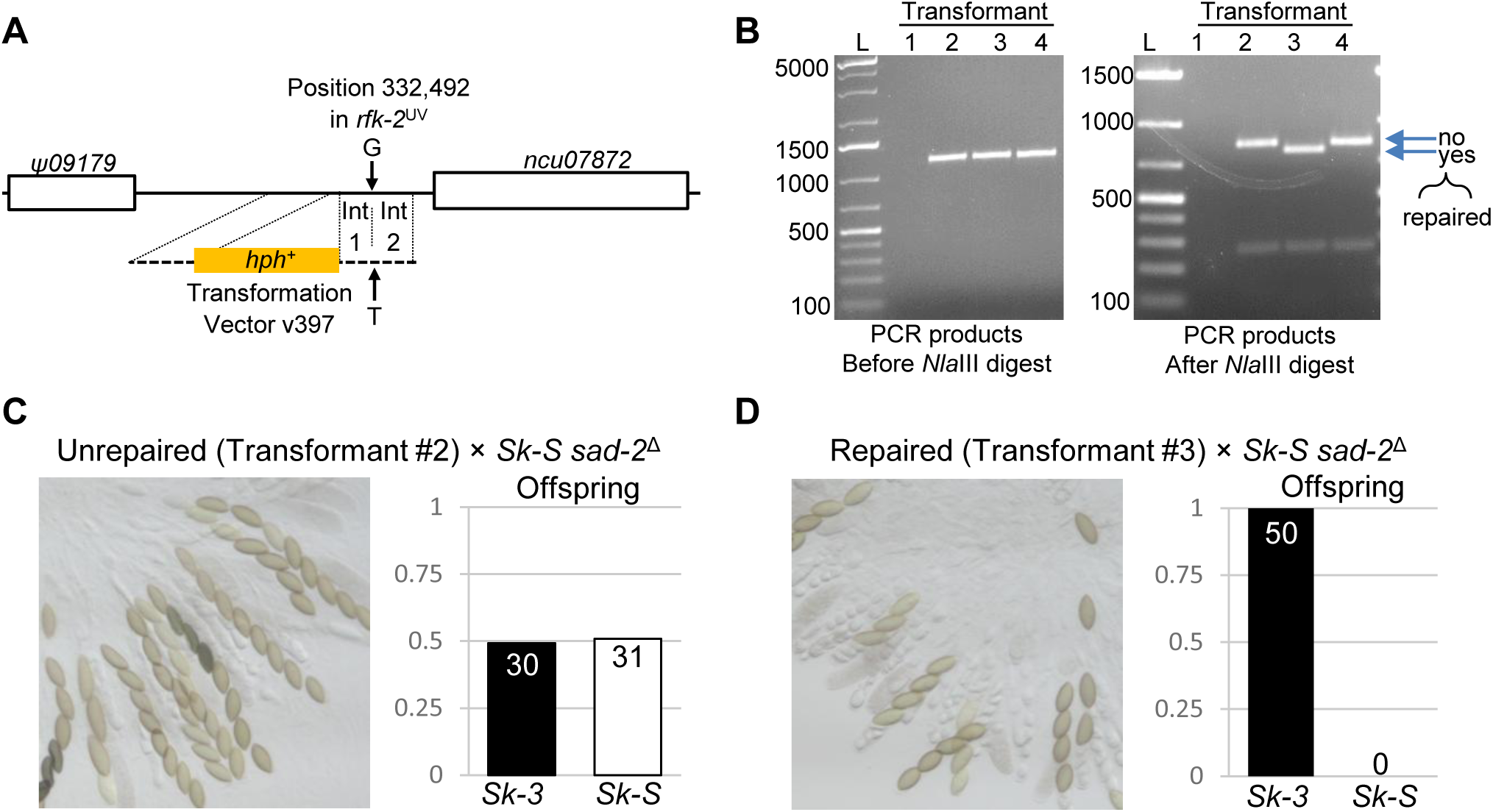
Repairing the mutation at Position 332,492 in an *rfk-2*^UV^ strain restores spore killing. (A) The strategy for repairing Position 332,492 in an *rfk-2*^UV^ strain is diagrammed. The right flank of v397 overlaps Position 332,492. The *rfk-2*^UV^ transformation host ISU-4684 encodes a guanosine at Position 332,492, while the right flank in v397 carries a thymidine at this position. Recombination points in Interval 1 and Interval 2 are "not expected" and "expected", respectively, to change Position 332,492 from a guanosine to a thymidine. (B) DNA samples from four transformants were used as templates in PCR reactions with Primer Set 2415/2525. The PCR products were examined before and after restriction digestion with *Nla*III (5′-CATG-3′) by gel electrophoresis. The T > G transversion mutation eliminates a *Nla*III site. These results show that the T>G mutation in “Transformant #3” was repaired. Lanes are as follows: L, DNA Ladder; 1, Transformant 1 (ISU-5567, PCR failed); 2, Transformant 2, (ISU-5568); 3, Transformant 3 (ISU-5569); 4, Transformant 4 (ISU-5571). (C and D) Homokaryotic isolates were obtained from two of the transformants and examined in spore killing assays. (C) Asci from “Transformant 2” (ISU-5856) × *Sk-S sad-2*^Δ^ (ISU-5563) are shown. Quotes are used to indicate that “Transformant 2” is actually a homokaryotic isolate derived from Transformant 2. Offspring (n = 61) were collected and genotyped with respect to *Sk-3* and *Sk-S.* These data show that spore killing did not occur in the cross and *Sk-3* failed to drive. (D) “Transformant 3” (ISU-5859) × *Sk-S sad-2*^Δ^ (ISU-5563). Offspring (n = 50) were collected and genotyped as in Panel C. These data show that *Sk-3* was transmitted to all viable offspring through spore killing.

### The *rfk-2* locus

To demarcate the borders of the *rfk-2* gene, we deleted a series of DNA intervals from an *Sk-3 rfk-2^+^* strain and examined the effects of these deletions on spore killing. Interval *i388* is located centromere-distal of *i350* and spans a putative pseudogene denoted as ψ*09179* (Figure 4A). Interval *i358* is located centromere-proximal of *i350* and spans gene *ncu07872*. Intervals *i397* and *i383* are found within *i350*. We found that deletions of *i388*, *i397*, and *i358* had no effect on spore killing, while deletion of *i383* eliminated it (Figure 4, B–D). The positions of the intervals and their deletion phenotypes reveal that the *rfk-2* gene must lie between intervals *i397* and *i358*. We refer to this 1039 bp region as the *rfk-2* locus.

**Figure 4.**
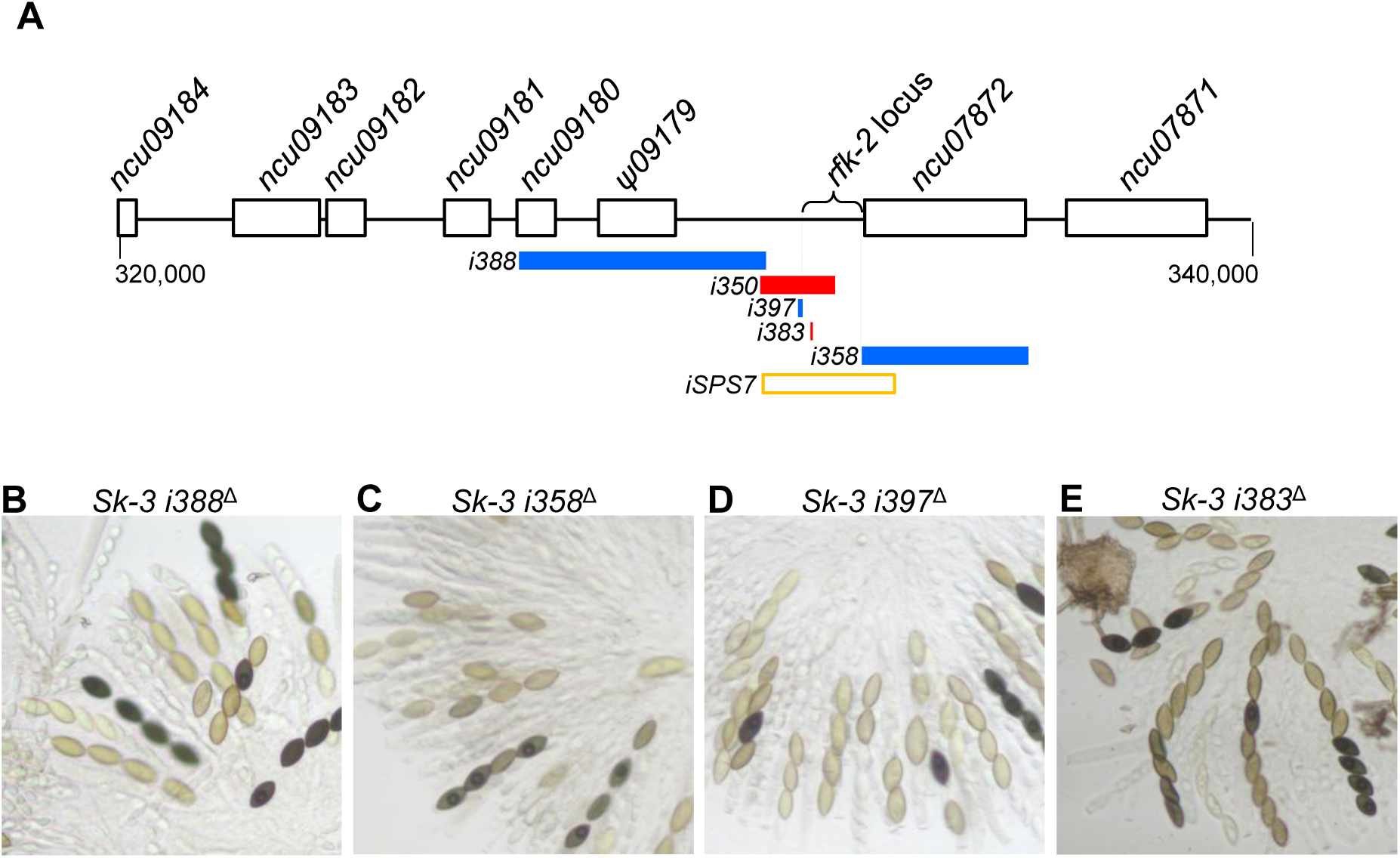
Gene deletion analysis of the *rfk-2* locus. (A) A map of Positions 320,000 through 340,000 of Chromosome III in the *Sk-3* reference sequence is shown. This map is the same as the one provided in Figure 2A, except here it has been annotated to show the locations of six intervals that were investigated with deletion assays (*i350, i358, i383, i388, i397*) or incorporated into an *rfk-2^+^* transgene (*iSPS7*). Red bars are used for intervals that result in loss of spore killing upon deletion. Blue bars are used for intervals that have no effect on spore killing upon deletion. The open rectangle with orange border (*iSPS7*) marks the *rfk-2*^+^-spanning interval that was transferred to *Sk-S* strains. (B–E) Asci from crosses between deletion strains (as males) with an *Sk-S sad-2*^Δ^ strain (as female) are shown. Male genotypes are indicated above each image. These results show that the critical sequences for spore killing are located between *i397* and *i358*. The strain names of the parents used in each cross are as follows: B) ISU-5295 × ISU-3036; C) ISU-5434 × ISU-3037; D) ISU-5462 × ISU-3036; and E) ISU-5211 × ISU-3037.

### *rfk-2^+^* is insufficient for spore killing in *Sk-S* × *Sk-S* crosses

To test if *rfk-2^+^* is sufficient for spore killing, we transferred the *rfk-2* locus to a common transgene insertion location on Chromosome I in an *Sk-S* strain and examined the resulting *Sk-S rfk-2^+^* strain in crosses. However, we did not detect abnormalities in asci from a cross between *Sk-S rfk-2+* and *Sk-S* mating partners (Figure S1). This finding suggested that *rfk-2^+^* is insufficient for spore killing in an *Sk-S* genetic background. Next, to determine if ectopic expression of *rfk-2^+^* could complement an *rfk-2*-deficient strain, we crossed *Sk-S rfk-2^+^* with *Sk-3 i383*^Δ^ (recall that deletion of *i383* eliminates spore killing) and isolated offspring with various genotypes for examination in spore killing assays. The *Sk-S rfk-2^+^*genotype produced phenotypically normal asci and its *rfk-2^+^*transgene was transmitted to offspring at a rate of 0.57 (Figure 5, A and D; Chi square, p = 0.28). Similarly, the *Sk-3 i383*^Δ^ genotype produced phenotypically normal asci, and *Sk-3* was transmitted to offspring at a rate of 0.46 (Figure 6, B and E; Chi square, p = 0.55). In contrast to the other two examined genotypes, the *Sk-3 i383*^Δ^ *rfk-2^+^* genotype produced asci with a spore killing phenotype and *Sk-3 i383*^Δ^ was transmitted to offspring at a rate of 1.0 (Figure 5, C, F, and G, Chi square, p < 0.0001). This finding demonstrates that ectopic expression of *rfk-2^+^* restores spore killing and gene drive to a *Sk-3 i383*^Δ^ genotype. Interestingly, *rfk-2^+^* was found in only 57% of offspring from this last cross, indicating that while ectopic expression of *rfk-2^+^* can drive *Sk-3 i383*^Δ^ through sexual reproduction it cannot do the same for itself (Figure 5G, Chi square, p = 0.51).

**Figure 5.**
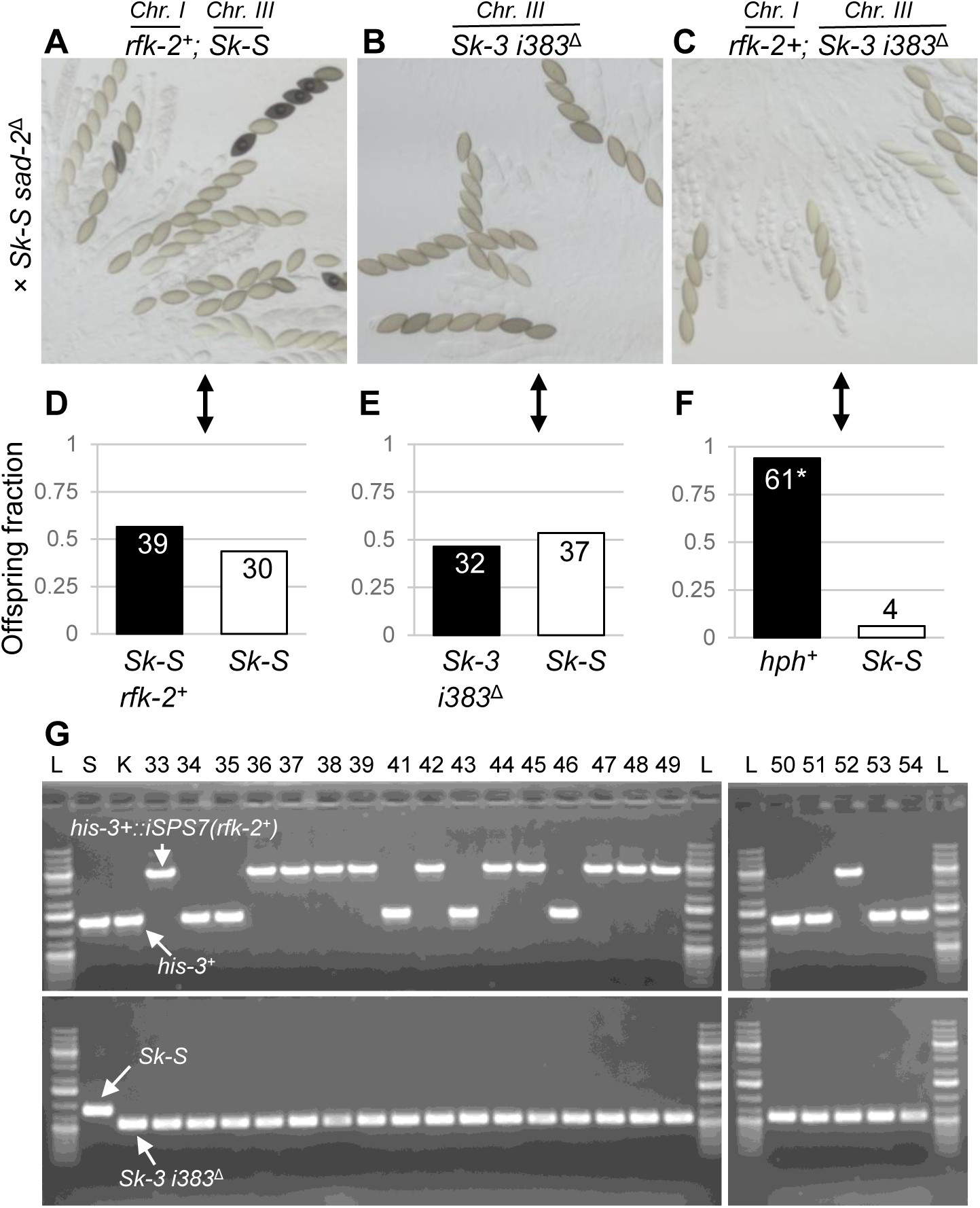
*rfk-2+* restores spore killing to *Sk-3 i383*^Δ^ *in trans*. (A–C) Asci from unidirectional crosses are shown. *Sk-S sad-2*^Δ^ was used as female and (A) *Sk-S rfk-2^+^*, (B) *Sk-3 i383*Δ, and (C) *Sk-3 i383*^Δ^ *rfk-2+* were used as males. Spore killing was only observed when *Sk-3 i383*^Δ^ *rfk-2^+^* was used as male (Panel C). (D) Offspring (n = 69) from the cross depicted in Panel A were genotyped for *rfk-2^+^*. *rfk-2^+^* was transmitted to offspring according to Mendelian inheritance (Chi square, p = 0.28). (E) Offspring (n = 69) from the cross depicted in Panel B were genotyped for *Sk-3 i383*^Δ^. *Sk-3 i383*^Δ^ was transmitted to offspring according to Mendelian inheritance (Chi square, p = 0.55). (F) Offspring (n = 65) from the cross depicted in Panel C were genotyped for resistance to hygromycin, which indicates the presence of *Sk-3 i383*^Δ^ and/or *rfk-2^+^.* Nearly all offspring were resistant to hygromycin. (G) Approximately 1/3 (n = 21) of the hygromycin resistant offspring from the chart in “F” were genotyped by PCR. All 21 (100%) had the *Sk-3 i383*^Δ^ genotype, and 12 of 21 also carried *rfk-2^+^*. These data demonstrate *Sk-3 i383*^Δ^ was transmitted to all 21 of the hygromycin-resistant offspring against the rules of Mendelian inheritance (Chi square, p < 0.0001), while *rfk-2^+^* was transmitted according to Mendelian inheritance (Chi square, p = 0.51). The following parents were used in the crosses: (A and D) ISU-5846 × ISU-5563, (B and E) ISU-5848 × ISU-5564, (C, F, and G) ISU-5852 × ISU-5563.

**Figure 6.**
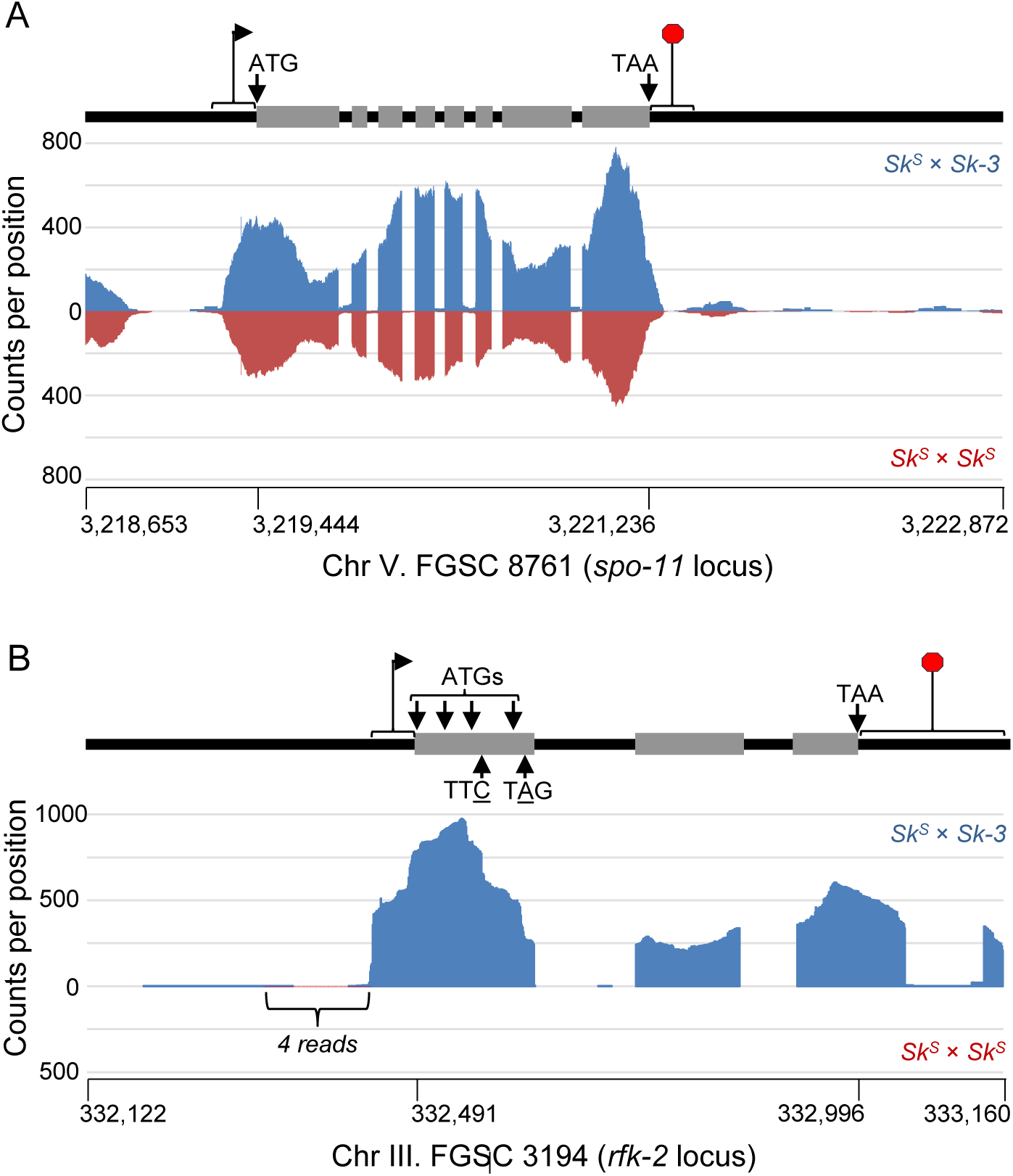
*rfk-2* transcript detection. (A) RNA sequencing reads corresponding to *Sk-S* × *Sk-3* (blue) and *Sk-S* × *Sk-S* (red) crosses were mapped to the genome of an *N. intermedia Sk-S* strain. Only reads aligning to the *spo-11* locus on Chromosome V are examined in the chart. This chart demonstrates that read counts across the meiotic endonuclease *spo-11* are similar between the two datasets, and thus the datasets are likely of similar quality with respect to meiotic gene expression. The x-axis indicates position along the *spo-11* locus. The y-axis indicates total read count at each position. A map of *spo-11* is shown above the chart. Gray bars mark the locations of predicted coding sequences. Transcription start site (bracket and bent arrow) and stop site (bracket and stop sign) are approximations. (B) Reads that did not map to the *N. intermedia Sk-S* genome were mapped to the *rfk-2* locus (Chromosome III, Positions 332,122-333,160) of the *Sk-3* reference genome. Only four reads from the *Sk-S* × *Sk-S* dataset mapped to this location. A model of *rfk-2* features is shown in the map above the chart. Features are as follows: An approximate location of the transcription start site (bracket and bent arrow); candidate start codons (ATGs); a codon that undergoes C to T editing (TT<u>C</u>); a codon that undergoes A to G/I editing (T<u>A</u>G); three exons with coding sequences (gray bars); two introns between the predicted coding exons (black bars); and an approximate location of the 3′-UTR/termination signal. Position 332,492, which is mutated in *rfk-2*^UV^, encodes the second nucleotide of the first candidate start codon. The third intron in the diagram is located in the *rfk-2* 3’ UTR and/or the 5’ UTR of the flanking gene *ncu07872*.

### *rfk-2* transcripts undergo RNA editing

Our demonstration that *rfk-2^+^* restores spore killing and gene drive to an *Sk-3 i383*^Δ^ genotype from an ectopic location suggests that *rfk-2* encodes a protein that acts *in trans*. If true, then *rfk-2* mRNA should be present in perithecial cultures of *Sk-3* × *Sk-S* crosses. To test this hypothesis, RNA sequencing datasets of poly(A)-enriched RNA from *Sk-3* × *Sk-S* and *Sk-S* × *Sk-S* crosses were downloaded from NCBI’s SRA database. To confirm that the datasets were of similar quality, reads were first mapped to an *Sk-S* reference genome, and reads that mapped to the meiotic crossover initiation gene, *spo-11* (Rhoades et al. 2021), were examined. Read counts at each position across *spo-11* were similar between the datasets (Figure 6A), suggesting that the datasets to be of similar quality.

We next mapped reads to the *rfk-2* locus. We found that four reads from the *Sk-S* × *Sk-S* dataset and 1800 reads from the *Sk-3* × *Sk-S* dataset mapped to *rfk-2* (Figure 6B). The alignments suggest the presence of three introns, at least two of which are found within the predicted *rfk-2* coding region. The third intron appears to belong to the 5′ UTR of gene *ncu07872*. Interestingly, visual inspection of the read sequences suggested that one nucleotide in *rfk-2* transcripts undergoes a cytidine to uridine edit and a second nucleotide undergoes an adenosine to guanosine/inosine edit. To confirm these observations, we used RT-PCR in attempts to amplify a partial *rfk-2* cDNA from *Sk-3*× *Sk-S* and *Sk-S* × *Sk-S* perithecia. As expected, we were only able to amplify a cDNA from *Sk-3* × *Sk-S* perithecia (Figure 7, A and B). Sequencing of eight cDNA clones found that six of the eight clones originated from RNA templates that had been edited at both the C-to-U and A-to-G/I positions, while two originated from templates that had not been edited at either position (Figure 7C).

**Figure 7.**
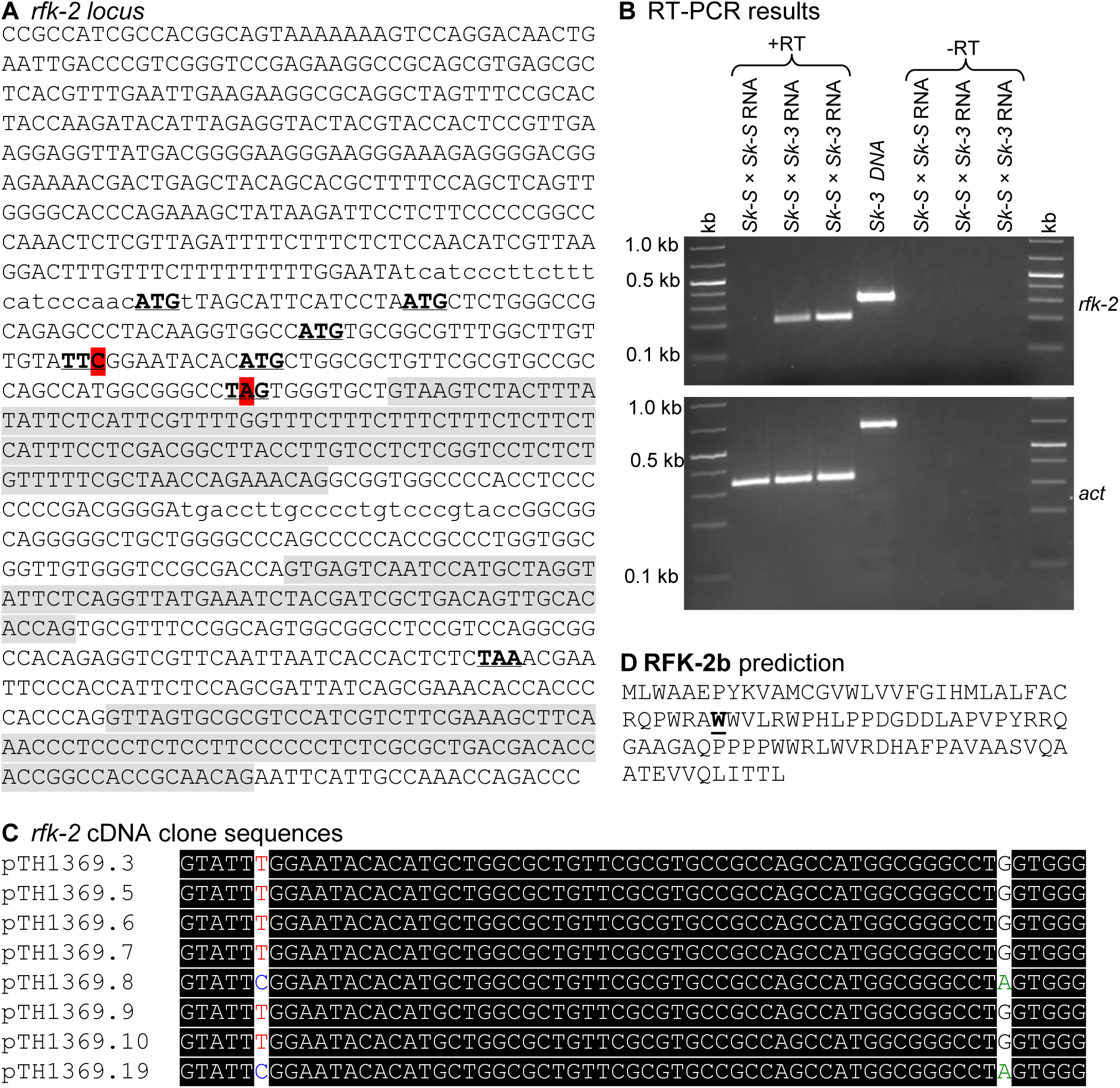
*rfk-2* cDNA isolation and sequencing confirms RNA editing of *rfk-2* transcripts. (A) The sequence of the *rfk-2* locus is shown. Four candidate ATG start codons are indicated with bold and underlined font, as is a TTC-phenylalanine codon that undergoes editing to a TTT-phenylalanine codon. The edited base of the TTC codon is highlighted in red. A TAG stop codon that undergoes editing to TGG-tryptophan codon is also indicated with bold and underlined font, with the edited base highlighted in red. Introns, as predicted from RNA sequencing data, are indicated with gray shading. The primer binding sites used for amplification of a partial cDNA are indicated by lowercase font. (B) RNA was isolated from unidirectional crosses between *Sk-S* (ISU-5564) × *Sk-S* (P6-07) and *Sk-S* (ISU-5564) × *Sk-3* (ISU-3291) perithecia 8 days post fertilization and used as template in RT-PCR reactions with Primer Set 2365/2468 for *rfk-2* and Primer Set 147/148 for *actin.* Reverse transcriptase was omitted (-RT) from some reactions as a control for DNA contamination of RNA samples. Of the two *Sk-S* × *Sk-3* samples, one was treated with DNAse I and purified with a column-based purification kit. This additional step appears to have been unnecessary as no genomic DNA contamination was detected in the samples not treated with DNAse I. (C) The *rfk-2* cDNA was cloned into pJET1.2 using the CloneJET PCR cloning kit and eight clones were sent for whole plasmid sequencing (Plasmidsaurus Inc.). The eight clones were identical except for the two positions that were predicted to undergo RNA editing according to the RNA sequencing dataset analysis. Specifically, six clones contained the edited base at both positions, while two clones contained neither edit. (D) Predicted RFK-2 protein sequence after editing of the UAG-stop codon to a UGG-tryptophan codon (the resulting tryptophan residue is indicted with bold and underlined font).

**Figure 8.**
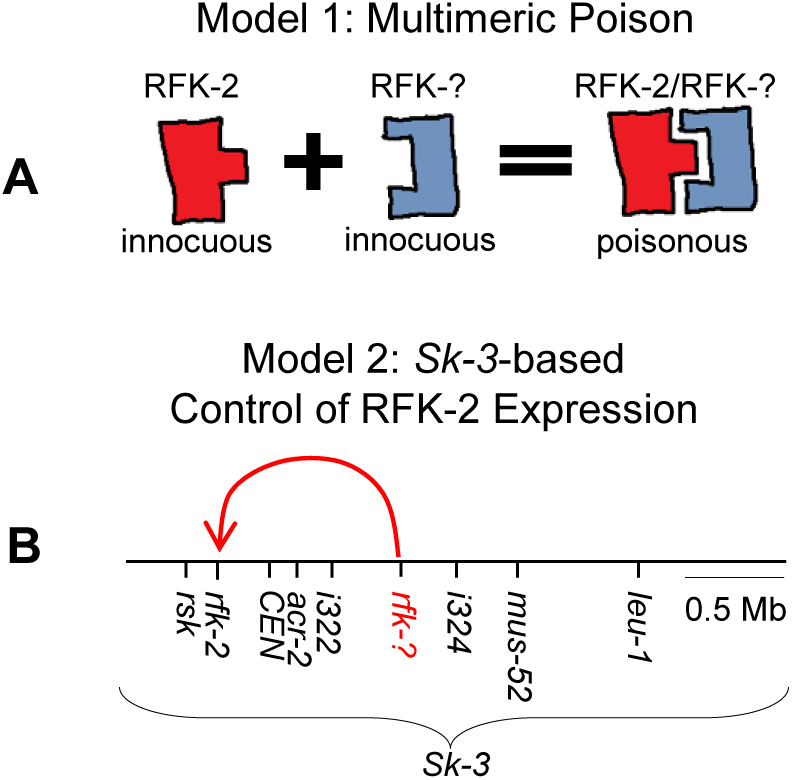
RFK-2 models. (A) RFK-2 may form part of a poisonous, multimeric compound. Each component of the multimeric compound is harmless in the absence of the other component. (B) RFK-2 alone may be poisonous, but its expression may require other genes within the *Sk-3* element. These genes could influence transcription rates and/or editing of *rfk-2* transcripts. A hypothetical gene "*rfk-?*", is depicted in red font. The red curved arrow is used to denote the influence of "*rfk-?*" on *rfk-2* expression. The depicted location of "*rfk-?*" between *i322* and *i324* was chosen for explanatory purposes only.

The above observations support a gene model where the *rfk-2* coding region begins with one of four possible AUG start codons (Figure 7A, underlined ATG codons) and contains two stop codons (an early UAG stop codon and a late UAA stop codon). The early UAG stop codon undergoes RNA editing to a UGG/UIG tryptophan codon. Before editing of the UAG stop codon, translation should produce a 13–36 amino acid protein (RFK-2a; the 36 amino acid length assumes the first of four candidate start codons is the actual start codon). After UAG editing, translation should produce a 78–101 amino acid protein (RFK-2b, Figure 7D). A search of NCBI’s ClusteredNR protein database (Sayers et al. 2019) with blastP and RFK-2b as query did not identify similar proteins (Altschul et al. 1997; Altschul et al. 2005). Additionally, a search of NCBI’s conserved domain database (Wang et al. 2023) with the same sequence as query failed to identify conserved domains. The lack of similar protein sequences and conserved domains in two of the most comprehensive collections of protein sequences and protein domains suggests that RFK-2 mediates spore killing through a novel mechanism.

## Discussion

Neurospora *Sk-3* is is thought to achieve gene drive through a poison-antidote system. While the antidote gene, *rsk*, has been known for over a decade, the identity of the poison gene has remained elusive. In this study, we have shown that a gene called *rfk-2* is required by *Sk-3* to kill spores. Below we discuss the possibility that *rfk-2* is *Sk-3*’s poison gene.

Our discovery that *rfk-2*^UV^ is allelic with DNA Interval *i350* was somewhat fortuitous. We sequenced a collection of *rfk-2*^UV^ strains to identify a set of SNVs that could then be used as markers to refine the location of *rfk-2*^UV^-through recombination mapping. Because we had previously mapped *rfk-2*^UV^ to the left of *i322* on Chromosome III, we focused on identifying SNVs in this region. In total, we identified 41 *rfk-2*^UV^-specific SNVs (Table S3). However, replacing *i350* with *hph* in an *Sk-3 rfk-2*^+^ strain disrupted spore killing, and our subsequent finding that *i350* spans the causative mutation in *rfk-2*^UV^ made further refinement of *rfk-2*’s location through mapping of recombination points unnecessary.

To demarcate the borders of *rfk-2*, we performed a series of interval deletion assays by building on our findings with *i350*. The five interval deletions presented in this study represent a subset of a much larger set of interval deletions that were performed to demarcate the borders of *rfk-2*. In total, and with the help of 54 undergraduate research assistants over three years, we deleted 37 different intervals (Figure S2). Seven are located centromere-distal to *i350*, 19 overlap or are found within *i350*, and 10 are found centromere-distal of *i350* (Figure S2). Fourteen undergraduate students have completed Senior Theses on the effects of one or more of these interval deletions on spore killing, and their theses are available for review at Illinois State University’s Research and eData repository (Bowen 2024; Krivograd 2024; Lee 2024; Marcheschi 2024; Munn 2024; Okleiteris 2024; Patel 2024; Damkoehler 2025; Klann 2025; Paulikas 2025; Rimmey 2025; Bhatt 2026; Spriggs 2026; Tobin 2026).

Neurospora *Sk-2* and *Sk-3* appear to share a common evolutionary origin. For example, *Sk-2* and *Sk-3* both use alleles of the *rsk* gene to provide resistance to spore killing and they both span millions of base pairs of the same region of Chromosome III (Hammond et al. 2012; Svedberg et al. 2018). Despite their similarities, their poison genes appear to be unique. The *Sk-2* killer protein is encoded by the *rfk-1* gene, which encodes two variants called RFK-1a and RFK-1b (Rhoades et al. 2019). RFK-1a is a 102 amino acid protein that is produced when translation of *rfk-1* transcripts stops at an early UAG stop codon. In sexual tissue, the early stop codon is edited to UIG (I is Inosine) and translation produces the 130 amino acid RFK-1b. Interestingly, transferring a 1,481 bp *rfk-1^+^*-spanning DNA interval called *AH36* to an *Sk-S* genetic background results in a strain that produces mostly aborted asci when crossed with an *Sk-S sad-2*^Δ^ mating partner (Rhoades et al. 2019). Current models hold that the RFK-1 poison is produced within asci of these crosses, and because *Sk-S* genetic backgrounds lack a compatible antidote, most asci succumb to the effects of the poison.

Although the evolutionary relationship between *rfk-1* (in *Sk-2*) and *rfk-2* (in *Sk-3*) is unclear, there are similarities in how the genes function. For example, both genes produce transcripts that undergo RNA editing. Additionally, both genes are required by their respective host elements to kill spores. A third similarity exists in the presence of early and late stop codons in transcripts from both genes, with the early UAG stop codon in both *rfk-1* and *rfk-2* transcripts undergoing editing to a UIG codon in sexual tissue. Two minor caveats to this last point are that we have not specifically examined vegetative tissue to confirm that *rfk-2* transcripts are edited only during the sexual cycle (although others have shown that A-to-I mRNA editing is sexual tissue specific in Neurospora fungi (Liu et al. 2017)), and we have not tested our current model of *rfk-2*’s translational reading frame.

Given the similarities between *rfk-1* and *rfk-2*, we were surprised to find that *rfk-2^+^* does not trigger ascus abortion when present in an *Sk-S* homozygous cross. This led us to believe that something might be wrong with the *rfk-2*^+^ transgene, a hypothesis that we discarded after finding that the same transgene restores spore killing *in trans* to an *Sk-3 i383*^Δ^ genotype. These findings suggest that *rfk-2* requires at least one other *Sk-3*-encoded genetic element to mediate spore killing. One possibility is that RFK-2 interacts with a second *Sk-3* protein to form a multimeric killer (Figure 9, Model 1). A second possibility is that RFK-2 alone is the killer, but other genetic elements within *Sk-3* control *rfk-2* transcription rates and/or post-transcriptional modifications (Figure 9, Model 2).

Perhaps one of the most fascinating aspects of *Sk-2* and *Sk-3* is the possibility that these elements have descended from a common ancestral Spore killer. Their presence in the same genomic region and their shared use of *rsk* as their resistance protein supports a shared evolutionary history, but the differences between their *required for killing* genes, so far, suggests that their evolutionary histories may be as complex as their transmission mechanisms.

## DATA AVAILABILITY

The authors affirm that all data necessary for confirming the conclusions of the article are present within the article, figures, tables, and publicly available datasets (NCBI BioProject PRJNA1490908). Strains and plasmids are available upon request.

## ACKNOWLEDGEMENTS

This study was made possible by the pioneering work of Barbara Turner, Namboori Raju, David Perkins, and others. We thank the Fungal Genetics Stock Center for serving as a repository for the Spore killer strains (McCluskey et al. 2010) and members of the Hammond Laboratory for technical assistance. TH conceived the project, designed the experiments, provided technical assistance, analyzed the results, and wrote the manuscript. Major contributions to the experiments presented in each figure are as follows: Figure 1 (NR, JL, DB, TH); Figure 2 (SPS, TH), Figure 3 (SPS, SM, TH), Figure 4 (SPS, NR, TH, undergraduate assistants); Figure 5 (SPS, PJ, TH); Figure 6 (SPS, SM, PJ, TH); Figure 7 (TH); Figure 8 (JS, TH); Figure 9 (TH). We thank Jesper Svedberg for reviewing the bioinformatics data and providing helpful comments on the *rfk-2* gene model. All authors reviewed and approved the final version of the manuscript. This work was supported by an award to TH from the National Science Foundation (2005295). This work was also in part supported by the U.S. Department of Agriculture, Agricultural Research Service. Disclaimer: Mention of trade names or commercial products in this publication is solely for the purpose of providing specific information and does not imply recommendation or endorsement by the US Department of Agriculture. USDA is an equal opportunity provider and employer.

We also thank the following Illinois State University undergraduate research assistants for performing interval deletions, spore killing assays, and gene drive assays. The students and the intervals they studied are as follows: Alexander Davis (Aug 2024–May 2025, *i399*), Alexandra Spriggs (Aug 2024–May 2026, *i401*, Senior Thesis), Allison Rimmey (Aug 2024– May 2026, *i358*, Senior Thesis), Anita Kovalaske (Aug 2025–t.b.d., *i395*), Brandon Green (Aug 2023–May 2024, *i390*), Bella Rohrig (Aug 2023–Dec 2024, *i388* and *i397*), Bethany Wilder (Jan 2023–May 2024, *i350*), Carolina Okleiteris (Aug 2022–May 2024, *i377*, Senior Thesis), Colin Miller (Aug 2025–t.b.d., *i405*), Daniella Aguado (Jan 2025–May 2025, *i407*), David Liu (Jan 2024–May 2025, *i386* and *i406*), Dayton Oblinger–Hammond (Aug 2023–May 2025, *i380* and *i407*), Diana Segerstrom (Aug 2023–Dec 2023, *i386*), Dylan Beilfuss (Jan 2026–May 2026, *i398*), Ellana Tobin (Aug 2024–t.b.d., *i402*, Senior Thesis), Emily Wooldridge (Jan 2025–May 2025, *i394*), Emma Schmidt (Aug 2025–t.b.d., *i403*), Erika Wells (Aug 2025–t.b.d., *i406*), Ethan Howard (Jan 2025–May 2025, *i397*), Grayson Mirabile (Jan 2026–May 2026, *i399*), Jack Deshand (Aug 2025–May 2026, *i382*), Jalen Lee (Aug 2022–May 2024, *i384*, Senior Thesis), Jenna McAleese (Aug 2025–May 2026, *i397*), John Munn (Aug 2022–May 2024, *i376*, Senior Thesis), Josh Stearns (Aug 2025–t.b.d., *i400*), Julia Sands (Aug 2023–May 2025, *i385* and *i405*), Kenzie Berns (Jan 2026–t.b.d., *i406*), Kole Damkoehler (Jan 2024–May 2025, *i386* and *i408*, Senior Thesis), Lana Seyl (Jan 2026–t.b.d., *i399*), Lauren Wilson (Jan 2026–May 2026, *i398*), Layla Jones (Aug 2023–May 2024, *i350*), Lydia Grampp (Jan 2024–May 2024, *i350*), Maddie Lynn (Jan 2025–May 2024, *i391*), Makenna Klann (Aug 2023–May 2025, *i382* and *i400*, Senior Thesis), Megan Haynes (Aug 2025–May 2026, *i383*), Michael Marcheschi (Aug 2022–May 2024, *i375*, Senior Thesis), Mollie Paeth (Aug 2024–May 2025, *i395*), Natalia Rojas Cruz (Aug 2023–Dec 2023, *i350*), Nevaeh McComb (Jan 2025–May 2025, *i358*), Nevyn Laframboise (Aug 2025–May 2026, *i396*), Osprey Baughman (Aug 2025–t.b.d., *i399*), Paulina Paulikas (Aug 2023–May 2025, *i383* and *i394*, Senior Thesis), Pedro Miguel Galvan (May 2022–May 2024, *i389*), Princy Patel (Aug 2022–May 2024, *i378*, Senior Thesis), Reaghan Williams (Aug 2024– May 2026, *i404*), Reece Bohms (Jan 2025–May 2025, *i397*), Sarah Martin (Jan 2025–May 2025, *i396*), Sean Gingrich (Aug 2023–Dec 2023, *i391*), Shadiyat Batula (Aug 2022–May 2025, *i372* and *i403*), Sophie Krivograd (Aug 2022–May 2024, *i374*, Senior Thesis), Tapasvi Bhatt (Aug 2024–May 2026, *i398*, Senior Thesis), Thera Bowen (Aug 2022–May 2024, *i373*, Senior Thesis), Tim Kedzierzawski (Aug 2024–May 2025, *i396*), Uzay Togay (Aug 2025–t.b.d., *i407*), Valerie Padula (Aug 2025–t.b.d., *i408*), Valerie Quevreaux (Jan 2025–May 2025, *i402*), Zachary Benbow (Aug 2023–May 2024, *i381*).

## LITERATURE CITED

Altschul SF et al. 1997. Gapped BLAST and PSI-BLAST: a new generation of protein database search programs. Nucleic Acids Res. 25(17):3389–3402. 10.1093/nar/25.17.3389

Altschul SF et al. 2005. Protein database searches using compositionally adjusted substitution matrices. FEBS J. 272(20):5101–5109. 10.1111/j.1742-4658.2005.04945.x

Basenko EY et al. 2018. FungiDB: An integrated bioinformatic resource for fungi and oomycetes. J Fungi. 4(1):39. 10.3390/jof4010039

Bhatt T. 2026. Qualitative and quantitative analysis of the *Sk-3 i398* interval in *Neurospora crassa*. Sr Theses – Biol Sci. 13. https://ir.library.illinoisstate.edu/stbs/13. 10.30707/1783431597.456829

Bowen T. 2024. Refining the *rfk-2* locus of *Sk-3* using the DNA Interval v373 in *Neurospora crassa*. Sr Theses – Biol Sci. 4. https://ir.library.illinoisstate.edu/stbs/4. 10.30707/1734473261.414558

Campbell JL, Turner BC. 1987. Recombination block in the Spore killer region of Neurospora. Genome Natl Res Counc Can. 29(1):129–135.

Damkoehler K. 2025. Determining the effects that deletion of *i386* and *i408* have on *Sk-3*-type spore killing. Sr Theses – Biol Sci. 10. https://ir.library.illinoisstate.edu/stbs/10. 10.30707/1764607194.332424

Dawe RK. 2022. The maize abnormal chromosome 10 meiotic drive haplotype: a review. Chromosome Res. 30(2–3):205–216. 10.1007/s10577-022-09693-6

Desjardins AE et al. 2004. Deletion and complementation of the mating type (MAT) locus of the wheat head blight pathogen *Gibberella zeae*. Appl Environ Microbiol. 70(4):2437–2444. 10.1128/AEM.70.4.2437-2444.2004

Freitag M, Williams RL, Kothe GO, Selker EU. 2002. A cytosine methyltransferase homologue is essential for repeat-induced point mutation in *Neurospora crassa*. Proc Natl Acad Sci. 99(13):8802–8807. 10.1073/pnas.132212899

Galagan JE et al. 2003. The genome sequence of the filamentous fungus *Neurospora crassa*. Nature. 422(6934):859–868. 10.1038/nature01554

Hall TA. 1999. BioEdit: a user-friendly biological sequence alignment editor and analysis program for Windows 95/98/NT. Nucleic Acid Symp Ser. 41:95–98

Hammond TM, Rehard DG, Xiao H, Shiu PKT. 2012. Molecular dissection of Neurospora Spore killer meiotic drive elements. Proc Natl Acad Sci U S A. 109(30):12093–12098. 10.1073/pnas.1203267109

Harvey AM et al. 2014. A critical component of meiotic drive in Neurospora is located near a chromosome rearrangement. Genetics. 197(4):1165–74. 10.1534/genetics.114.167007

Kim D et al. 2019. Graph-based genome alignment and genotyping with HISAT2 and HISAT-genotype. Nat Biotechnol. 37(8):907–915. 10.1038/s41587-019-0201-4

Klann M. 2025. Examining the mechanism of spore sacs undergoing *Sk-3*-based spore killing after deletion of *Neurospora crassa* DNA intervals *i382* and *i400*. Sr Theses – Biol Sci. 9. https://ir.library.illinoisstate.edu/stbs/9. 10.30707/1764607637.096177

Krivograd S. 2024. Investigating *Sk-3*-based spore killing through DNA deletion in *Neurospora crassa*. Sr Theses – Biol Sci. 2. https://ir.library.illinoisstate.edu/stbs/2/ 10.30707/1734473255.348925

Langmead B, Salzberg SL. 2012. Fast gapped-read alignment with Bowtie 2. Nat Methods. 9(4):357–359. 10.1038/nmeth.1923

Larracuente AM, Presgraves DC. 2012. The selfish *Segregation distorter* gene complex of *Drosophila melanogaster*. Genetics. 192(1):33–53. 10.1534/genetics.112.141390

Lee J. 2024. Studying a spore killing gene in *Neurospora crassa*. Sr Theses – Biol Sci. 7. https://ir.library.illinoisstate.edu/stbs/7/ 10.30707/1734536653.457732

Li H et al. 2009. The Sequence Alignment/Map format and SAMtools. Bioinformatics. 25(16):2078–2079. 10.1093/bioinformatics/btp352

Liu H et al. 2017. A-to-I RNA editing is developmentally regulated and generally adaptive for sexual reproduction in Neurospora crassa. Proc Natl Acad Sci U S A. 114(37):E7756– E7765. 10.1073/pnas.1702591114

Lohmar JM et al. 2022. A-to-I mRNA editing controls spore death induced by a fungal meiotic drive gene in homologous and heterologous expression systems. Genetics. 221(1):iyac029. 10.1093/genetics/iyac029

Marcheschi M. 2024. Investigating the role of DNA interval *v375* in spore killing by *Neurospora crassa Sk-3*. Sr Theses – Biol Sci. 6. https://ir.library.illinoisstate.edu/stbs/6. 10.30707/1734473246.481567

Margolin BS, Freitag M, Selker EU. 1997. Improved plasmids for gene targeting at the *his-3* locus of *Neurospora crassa* by electroporation. Fungal Genet Newsl. 44:34–36

McCluskey K, Wiest A, Plamann M. 2010. The Fungal Genetics Stock Center: a repository for 50 years of fungal genetics research. J Biosci. 35(1):119–126

Munn J. 2024. Investigating the role of the DNA interval *v376* on *Sk-3* Spore killing in *Neurospora crassa*. Sr Theses – Biol Sci. https://ir.library.illinoisstate.edu/stbs/1. 10.30707/1734473241.700929

Ninomiya Y, Suzuki K, Ishii C, Inoue H. 2004. Highly efficient gene replacements in Neurospora strains deficient for nonhomologous end-joining. Proc Natl Acad Sci U S A. 101(33):12248–12253. 10.1073/pnas.0402780101

Okleiteris C. 2024. Investigating the role of DNA interval *v377* in spore killing by *Neurospora crassa Sk-3*. Sr Theses – Biol Sci. 5. https://ir.library.illinoisstate.edu/stbs/5. 10.30707/1734473236.364631

Patel P. 2024. Identifying the location of the *rfk-2* spore killer gene on Chromosome III of *Neurospora crassa*. Sr Theses – Biol Sci. 3. https://ir.library.illinoisstate.edu/stbs/3. 10.30707/1734473226.462803

Paulikas P. 2025. Investigating *Sk-3* based Spore killing in *Neurospora crassa* through deletion analysis of DNA Intervals *i383* and *i394*. Sr Theses – Biol Sci. 8. https://ir.library.illinoisstate.edu/stbs/8. 10.30707/1764607804.375978

Perkins DD, Radford A, Sachs MS. 2000. The Neurospora Compendium: Chromosomal Loci. Academic Press.

Raju NB. 1979. Cytogenetic behavior of spore killer genes in neurospora. Genetics. 93(3):607– 623.

Raju NB. 1980. Meiosis and ascospore genesis in Neurospora. Eur J Cell Biol. 23(1):208–223

Rhoades N et al. 2021. Recombination-independent recognition of DNA homology for meiotic silencing in Neurospora crassa. Proc Natl Acad Sci U S A. 118(33):e2108664118. 10.1073/pnas.2108664118

Rhoades NA et al. 2019. Identification of *rfk-1*, a meiotic driver undergoing RNA editing in Neurospora. Genetics. 212(1):93–110. 10.1534/genetics.119.302122

Rhoades NA, Webber EK, Hammond TM. 2020. A nonhomologous end-joining mutant for Neurospora sitophila research. Fungal Genet Rep. 64:Article 1. 10.4148/1941-4765.2172

Rimmey A. 2025. Investigating the effect of gene *ncu07872* deletion on Spore killing by *Neurospora Sk-3*. Sr Theses – Biol Sci. 11. https://ir.library.illinoisstate.edu/stbs/11/ 10.30707/1777915859.023576

Samarajeewa DA et al. 2014. Efficient detection of unpaired DNA requires a member of the Rad54-like family of homologous recombination proteins. Genetics. 198(3):895–904. 10.1534/genetics.114.168187

Sayers EW et al. 2019. Database resources of the National Center for Biotechnology Information. Nucleic Acids Res. 47(D1):D23–D28. 10.1093/nar/gky1069

Shiu PKT et al. 2006. SAD-2 is required for meiotic silencing by unpaired DNA and perinuclear localization of SAD-1 RNA-directed RNA polymerase. Proc Natl Acad Sci U S A. 103(7):2243–2248. 10.1073/pnas.0508896103

Smith Z, Bedore S, Spingler S, Hammond TM. 2016. A *mus-51* RIP allele for transformation of *Neurospora crassa*. Fungal Genet Rep. 62:1–7

Spriggs A. 2026. Investigating the role of *Sk-3* interval *i401* in gene drive by *Neurospora Spore Killer-3*. Sr Theses – Biol Sci. 12. https://ir.library.illinoisstate.edu/stbs/12. 10.30707/1783354831.720145

Svedberg J et al. 2018. Convergent evolution of complex genomic rearrangements in two fungal meiotic drive elements. Nat Commun. 9(1):4242. 10.1038/s41467-018-06562-x

Svedberg J et al. 2021. An introgressed gene causes meiotic drive in Neurospora sitophila. Proc Natl Acad Sci. 118(17) 10.1073/pnas.2026605118

The Galaxy Community. 2024. The Galaxy platform for accessible, reproducible, and collaborative data analyses: 2024 update. Nucleic Acids Res. 52(W1):W83–W94. 10.1093/nar/gkae410

Tobin E. 2026. Investigating the role of interval *i402* on *Sk-3*-based spore killing and gene drive in *Neurospora crassa*. Sr Theses – Biol Sci. 14. https://ir.library.illinoisstate.edu/stbs/14/ 10.30707/1784042475.274785

Turner BC, Perkins DD. 1979. *Spore killer*, a chromosomal factor in Neurospora that kills meiotic products not containing it. Genetics. 93(3):587–606

Velazquez A et al. 2022. Isolation of *rfk-2*^UV^, a mutation that blocks spore killing by *Neurospora Spore killer-3*. MicroPublication Biol. 2022. 10.17912/micropub.biology.000604

Vogel HJ. 1956. A convenient growth medium for Neurospora (Medium N). Microb Genet Bull. 13:42–43

Wang J et al. 2023. The conserved domain database in 2023. Nucleic Acids Res. 51(D1):D384– D388. 10.1093/nar/gkac1096

Westergaard M, Mitchell HK. 1947. Neurospora V. A Synthetic Medium Favoring Sexual Reproduction. Am J Bot. 34(10):573–577. 10.2307/2437339

Yu J-H et al. 2004. Double-joint PCR: a PCR-based molecular tool for gene manipulations in filamentous fungi. Fungal Genet Biol FG B. 41(11):973–981. 10.1016/j.fgb.2004.08.001

Zanders S, Johannesson H. 2021. Molecular mechanisms and evolutionary consequences of spore killers in ascomycetes. Microbiol Mol Biol Rev. 85(4):e0001621. 10.1128/MMBR.00016-21

