## Supplementary material for "Molecular characterization of *required for killing-2*: a gene required for spore killing by *Neurospora Sk-3*": Figure S1

**A** *Sk-S rfk-2*$^{+}$ × *Sk-S*

**B** *Sk-S rfk-2*$^{+}$ × *Sk-S sad-2*$^{\Delta}$

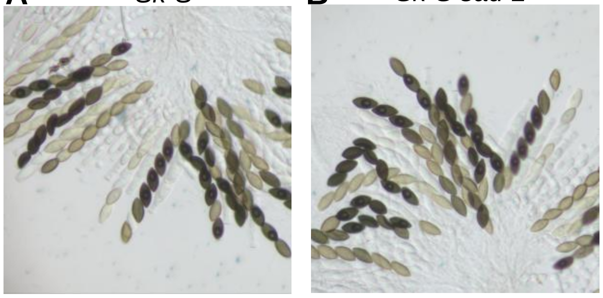

**Figure S1** Examination of *rfk-2*$^{+}$ in an *Sk-S* genetic background. A DNA interval called *iSPS7,* which spans the *rfk-2* locus, was cloned from an *Sk-3* strain and transferred to a standard transgene integration site downstream of *his-3* on Chromosome I in an *Sk-S* strain. The resulting *Sk-S rfk-2*$^{+}$ strain (ISU-5338) was then crossed as male with an *Sk-S* strain (F2-23) as female and an *Sk-S sad-2*$^{\Delta}$ strain (ISU-3036) as female. Spore killing asci were not detected in either cross, demonstrating that *rfk-2*$^{+}$ is insufficient for spore killing in an *Sk-S* genetic background.
