## Supplementary material for "Molecular characterization of *required for killing-2*: a gene required for spore killing by *Neurospora Sk-3*": Figure S2

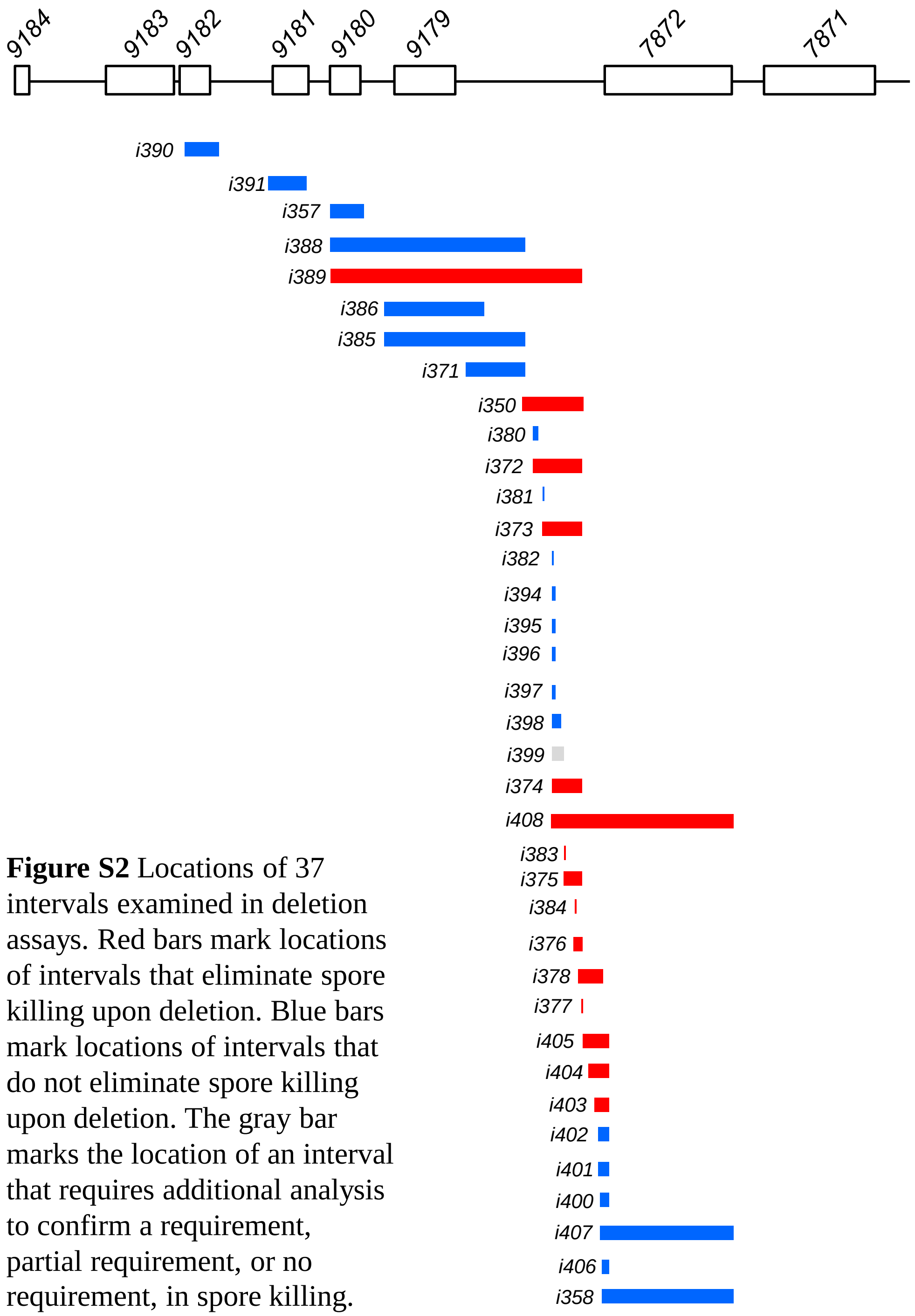


**Figure S2** Locations of 37 intervals examined in deletion assays. Red bars mark locations of intervals that eliminate spore killing upon deletion. Blue bars mark locations of intervals that do not eliminate spore killing upon deletion. The gray bar marks the location of an interval that requires additional analysis to confirm a requirement, partial requirement, or no requirement, in spore killing.
