## Supplementary material for "Molecular characterization of *required for killing-2*: a gene required for spore killing by *Neurospora Sk-3*": Table S1

**Table S1** Strain sources

| Strain name (alias) | Source |
|---|---|
| F2-23 (RTH1005.1) | Hammond et al. 2012 |
| F2-26 (RTH1005.2) | Hammond et al. 2012 |
| FGSC 10340 (RZS27.10) | Smith et al. 2016 |
| ISU-3036 (RTH1623.1) | Samarajeewa et al. 2014 |
| ISU-3037 (RTH1623.2) | Samarajeewa et al. 2014 |
| ISU-3291 (RDGR170.3) | Velazquez et al. 2022 |
| ISU-4677 (RTH0226.1.1) | Velazquez et al. 2022 |
| ISU-4683 (RNR323.7) | Velazquez et al. 2022 |
| ISU-4684 (RNR343.24) | Velazquez et al. 2022 |
| ISU-4773 (MAV214) | Velazquez et al. 2022 |
| ISU-5211 (RPPAK10.1) | F2-26 × Transformant of ISU-3291 with v383 |
| ISU-5295 (RIGR10.9) | F2-26 × Transformant of ISU-3291 with v388 |
| ISU-5331 (RNR343.5) | ISU-4683 × ISU-3291 |
| ISU-5332 (RNR343.6) | ISU-4683 × ISU-3291 |
| ISU-5333 (RNR343.15) | ISU-4683 × ISU-3291 |
| ISU-5334 (RNR343.17) | ISU-4683 × ISU-3291 |
| ISU-5335 (RNR343.19) | ISU-4683 × ISU-3291 |
| ISU-5337 (CNR427.2.1) | Transformant of ISU-3291 with v350 (Homokaryon isolated by screening single conidium-derived colonies) |
| ISU-5338 (CPJ3.3.3) | Transformant of FGSC 10340 with pSPS7.2 (Homokaryon isolated by screening single conidium-derived colonies) |
| ISU-5434 (RARim12.103) | F2-26 × Transformant of ISU-3291 with v358 |
| ISU-5462 (REHow10.12) | F2-26 × Transformant of ISU-3291 with v397 |
| ISU-5563 (RTH1925.13) | F2-23 × Transformant of FGSC-10340 with v417 |
| ISU-5564 (RTH1925.22) | F2-23 × Transformant of FGSC-10340 with v417 |
| ISU-5567 (TSPS28.1) | Heterokaryotic Transformant of ISU-4684 with v397 |
| ISU-5568 (TSPS28.2) | Heterokaryotic Transformant of ISU-4684 with v397 |
| ISU-5569 (TSPS28.3) | Heterokaryotic Transformant of ISU-4684 with v397 |
| ISU-5571 (TSPS28.4) | Heterokaryotic Transformant of ISU-4684 with v397 |
| ISU-5846 (RSPS23.12) | ISU-5211 × ISU-5338 |
| ISU-5848 (RSPS23.21) | ISU-5211 × ISU-5338 |
| ISU-5852 (RSPS23.11) | ISU-5211 × ISU-5338 |
| ISU-5856 (CSPS28.2.4) | Homokaryotic isolate from Heterokaryon ISU-5668 |

| | |
|---|---|
| ISU-5859 (CSPS28.3.2) | Homokaryotic isolate from Heterokaryon ISU-5669 |
| P6-07 | Hammond et al. 2012 |
