## Supplementary material for "Molecular characterization of *required for killing-2*: a gene required for spore killing by *Neurospora Sk-3*": Table S2

**TABLE S2** Primers used in this study

| NAME | SEQUENCE | |
|---|---|---|
| 12 (HPH-CEN-F) | AACTGATATTGAAGGAGCATTTTTTGG | *hph* CENTER |
| 13 (HPH-CEN-R) | AACTGGTTCCCGGTCGGCAT | *hph* CENTER |
| 147 (ACT-52113-A) | AAGAAGTTGCCGCCCTCGT | *actin* cDNA |
| 148 (ACT-52113-B) | CGGTTGGACTTGGGGTTGATG | *actin* cDNA |
| 2304 (v350-A) | TTGGCCCCATATCGTAACCTTCG | v350 LEFT |
| 2305 (v350-B) | AAAAAATGCTCCTTCAATATCAGTTGA CGCCACTCTCTGTCCCAACAA | v350 LEFT |
| 2306 (v350-C) | GAGTAGATGCCGACCGGGAACCAGTT CCTTGTCCTCTCGGTCCTCTCTGT | v350 RIGHT |
| 2307 (v350-D) | GGTAGCGTTCTGGGATGAGTGTGG | v350 RIGHT |
| 2308 (v350-E) | TGACGCCGGTTTAGCGTTGTAAG | v350 NESTED |
| 2309 (v350-F) | CGAGGGGACATGGACATTTGG | v350 NESTED |
| 2361 (v358-A) | GGCACCCAGAAAGCTATAAGATTCCT | v358 LEFT |
| 2362 (v358-B) | AAAAAATGCTCCTTCAATATCAGTTGG GTCTGGTTTGGCAATGAATTCTG | vV358 LEFT |
| 2363 (v358-C) | GAGTAGATGCCGACCGGGAACCAGTT ATAAATTGCTGGGCTAGGGAAGGTG | v358 RIGHT |
| 2364 (v358-D) | GGTGATGAATGGCGGATAGGTTCTT | v358RIGHT |
| 2365 (v358-E) | TCATCCCTTCTTTCATCCCAACATGT | *rfk-2* CDNA |
| 2365 (v358-E) | TCATCCCTTCTTTCATCCCAACATGT | v358 NESTED |
| 2366 (v358-F) | AGTTAGTTTGGCTCTGGATGACTGC | v358 NESTED |
| 2415 (HPH-TCU-FOR) | GGAGCGAGGCGATGTTCGGGGAT | T>G repair detection |
| 2435 (v375-A) | ATCACGGGGACTTGGCTCCATGC | v383 LEFT |
| 2436 (v375-B) | AAAAAATGCTCCTTCAATATCAGTTCC CTCTTTCCCTTCCCTTCCCCGT | v383 LEFT |

| | | |
|---|---|---|
| 2437 (v375-E) | ACCTCATGTCTCGGTGAAGGGCG | v383 NESTED |
| 2457 (v379)-CENF | TATTCGGCCTACCTGGCTGTGGC | v417 CENTER |
| 2458 (v379)-CENR | TAGTCGCGTGGAGCCAAGAGCGG | v417 CENTER |
| 2468 (v0382-F) | GGTACGGGACAGGGGCAAGGTCA | *rfk-2* cDNA |
| 2469 (v383-C) | GAGTAGATGCCGACCGGGAACCAGTTT GAGCTACAGCACGCTTTTCCAGC | v383 RIGHT |
| 2470 (v383-D) | CGATGGACGCGCACTAACCTGGG | v383 RIGHT |
| 2471 (v383-F) | CTGGGTGGGGTGGTGTTTCGCTG | v383 NESTED |
| 2507 (SK3-0524A) | AAAAGCGGCCGCGAGTGGCGTCTGCCT CGTGTTGA | *iSPS7*/rfk-2$^{+}$ transgene and T>G mutation detection |
| 2508 (SK3-0524B) | AAAAGCGGCCGCTCGGGAACTGGCGT CCTCTCGG | *iSPS7*/rfk-2$^{+}$ transgene and T>G mutation detection |
| 2520 (v394-A) | TCCAAAGGGAAGGACCGGGCACA | v397 LEFT |
| 2521 (v394-B) | AAAAAATGCTCCTTCAATATCAGTTCG TGAGCCGGAGCAGTCGTCGTA | v397 LEFT |
| 2523 (v394-D) | GACGCGCACTAACCTGGGTGGG | v397 RIGHT |
| 2524 (v394-E) | GGGACAGAGAGTGGCGTCTGCCT | v397 NESTED |
| 2525 (v394-F ) | CACTGCCGGAAACGCACTGGTGT | v397 NESTED and T>G repair detection |
| 2528 (v397-C) | GAGTAGATGCCGACCGGGAACCAGTT CCGCCATCGCCACGGCAGTAAAA | v397 RIGHT |
| 2608 (v417-A) | CGAGGTGTAGGCGTGGTGTGTTT | v417 LEFT |
| 2609 (v417-B) | GCCACAGCCAGGTAGGCCGAATAAGC GGGATGTCAAAGAAAGTTGCC | v417 LEFT |
| 2610 (v417-C) | CCGCTCTTGGCTCCACGCGACTACGCA GAACGCAAGTAAGAACGCA | v417 RIGHT |
| 2611 (v417-D) | TGACATCCAACACGAACGAGCACA | v417 RIGHT |
| 2612 (v417)-E | CTTGGATACCCGTGATGATTCTGATGA | v417 NESTED |
| 2613 (v417)-F | GCCCTCAGCCAACAATTTCTCCCT | v417 NESTED |
