## Supplementary material for "Molecular characterization of *required for killing-2*: a gene required for spore killing by *Neurospora Sk-3*": Table S3

**Table S2** SNV analysis. A total of 70 candidate SNVs were identified to the left of Interval *i322* among *six rfk-2*$^{UV}$ strains. An SNV is defined as a nucleotide that is different than that in the *Sk-3* reference sequence. Of the 70 candidate SNVs, 41 are unique to *rfk-2*$^{UV}$ while the remaining are shared by *rfk-2*$^{UV}$ and *rfk-2*$^{+}$. SNVs that are shared between *rfk-2*$^{UV}$ and *rfk-2*$^{+}$ may have been generated during the introgression of *Sk-3* from *N. intermedia* into *N. crassa* and/or during the routine subculturing of *N. crassa Sk-3* strains after introgression. Red font marks nculeotides that are different from the corresponding nucleotide in the reference sequence.

| Chr. III Position | FGSC 3194 nucleotide | *rfk-2*$^{UV}$ nucleotide | *rfk-2*$^{+}$ nucleotide |
|---|---|---|---|
| 76063 | G | A | A |
| 88338 | C | T | T |
| 93979 | C | T | T |
| 97864 | C | T | T |
| 97931 | C | T | T |
| 105609 | C | T | T |
| 105620 | C | T | T |
| 105638 | C | T | T |
| 124579 | C | T | T |
| 124612 | C | T | T |
| 129119 | G | A | A |
| 132286 | C | T | T |
| 134367 | C | T | T |
| 134369 | C | T | T |
| 332492 | T | G | T |
| 354271 | C | T | C |
| 371318 | C | G | G |
| 398579 | T | A | T |
| 458678 | G | A | G |
| 529071 | T | C | T |
| 546809 | G | A | A |
| 546819 | G | A | A |
| 557525 | C | T | C |
| 565421 | T | A | T |
| 569775 | A | G | A |
| 602515 | G | T | G |
| 610539 | A | G | A |
| 621595 | A | C | A |
| 643341 | G | A | G |
| 648013 | T | G | T |
| 666051 | T | C | T |

| | | | |
|---|---|---|---|
| 670128 | A | T | A |
| 676858 | T | C | T |
| 676862 | T | C | T |
| 698868 | A | G | A |
| 718303 | T | C | T |
| 734087 | T | G | T |
| 734089 | T | A | T |
| 734092 | A | T | A |
| 744137 | G | A | G |
| 754372 | C | T | T |
| 754384 | C | T | T |
| 756565 | C | T | T |
| 756577 | C | T | T |
| 756587 | C | T | T |
| 756687 | C | T | T |
| 756693 | C | T | T |
| 756730 | C | T | T |
| 756810 | C | T | T |
| 756906 | C | T | T |
| 758434 | A | G | A |
| 771671 | T | G | T |
| 789110 | T | G | T |
| 790996 | A | T | A |
| 799567 | T | C | T |
| 802086 | A | C | A |
| 807219 | A | G | A |
| 829870 | A | C | A |
| 833966 | T | G | T |
| 922912 | T | A | T |
| 929310 | T | A | T |
| 999988 | C | T | C |
| 1000972 | G | A | A |
| 1003885 | T | G | T |
| 1011687 | T | A | T |
| 1011694 | A | G | A |
| 1011695 | A | G | A |
| 1013944 | C | T | T |
| 1015430 | T | G | T |
| 1015444 | T | C | T |
